# Placebo treatment affects brain systems related to affective and cognitive processes, but not nociceptive pain

**DOI:** 10.1101/2023.09.21.558825

**Authors:** Rotem Botvinik-Nezer, Bogdan Petre, Marta Ceko, Martin A. Lindquist, Naomi P. Friedman, Tor D. Wager

**Affiliations:** Hebrew University of Jerusalem; Dartmouth College; University of Colorado, Boulder; Johns Hopkins University

## Abstract

Placebo analgesia is a replicable and well-studied phenomenon, yet it remains unclear to what degree it includes modulation of nociceptive processes. Some studies find effects consistent with nociceptive effects, but meta-analyses show that these effects are often small. We analyzed placebo analgesia in a large fMRI study (N = 392), including placebo effects on brain responses to noxious stimuli. Placebo treatment caused robust analgesia in both conditioned thermal and unconditioned mechanical pain. Placebo did not decrease fMRI activity in nociceptive pain regions, including the Neurologic Pain Signature (NPS) and pre-registered spinothalamic pathway regions, with strong support from Bayes Factor analyses. However, placebo treatment affected activity in pre-registered analyses of a second neuromarker, the Stimulus Intensity Independent Pain Signature (SIIPS), and several associated a priori brain regions related to motivation and value, in both thermal and mechanical pain. Individual differences in behavioral analgesia were correlated with neural changes in both thermal and mechanical pain. Our results indicate that processes related to affective and cognitive aspects of pain primarily drive placebo analgesia.

## Introduction

Throughout history, placebo effects have been variously considered as mysterious healing forces and tricks played upon the gullible by medical practitioners. Scientific research over the past decades has shown that placebo effects are neither of these. Rather, they are now understood to result from active, endogenous brain processes related to expectation, meaning, and predictive regulation of the body ^1–4^. A substantial part of the benefit of many kinds of treatments—including conventional drug therapies ^5–8^, surgery ^9,10^, acupuncture ^11,12^, psychotherapy ^13,14^, and more—is related to these psychological and brain processes. The study of placebo effects is thus the study of the internal brain processes that promote health and healing. It is imperative to understand these processes more completely in order to harness them in clinical care. In particular, it is still unknown at which level sensory experiences and physiological responses are affected by these internal processes.

Effective placebo treatments are thought to work by influencing ‘meaning-making’ systems ^15–18^– internal models of the world that shape our interpretations of sensory events, including their underlying causes and their implications for the future. These models determine our predispositions to react negatively or positively to events (e.g., attentional biases, evaluative biases, and mindsets ^19,20^), and facilitate enhanced or reduced reactions in perceptual and affective circuits ^17^. These internal models provide predictive signals that are integrated with incoming sensory experience to produce experienced bodily sensations and symptoms ^21–23^. For instance, current theories posit that pain is constructed by integrating ‘top-down’ context-based predictions with afferent nociceptive signals according to principles of Bayesian inference ^24,25^. Prediction errors (discrepancies between sensory input and predicted values) are propagated upstream for learning (refining the internal model). In this way, even placebo treatments– accompanied by suggestions, social cues, and prior experiences of success–can affect the neural construction of pain ^24,26–33^. Similar accounts have been increasingly used to explain symptoms and cognitive distortions in multiple disorders ^34–38^.

However, a critical open question remains: How fundamentally do placebos and other manipulations of predictive models affect sensory processing pathways? This question is important because its answer is the difference between a profound analgesic effect on responses (and plasticity) throughout the nervous system, effects on higher-level construction of pain experience, or transient biases in decision-making. By some accounts, predictions can propagate down multiple levels of perceptual hierarchies to affect sensory processing at the earliest stages– e.g., visual responses in visual thalamic pathways and V1 ^23,39^ and nociceptive processing in the spinal cord ^24^. In support of this view, early studies of placebo analgesia provided evidence for placebo or nocebo (aversive predictions) effects on the spinal cord ^40,41^, release of endogenous opioids ^42–46^, increases in putative descending pain-control pathways in the brainstem ^43,47,48^, and decreased pain-related activity in the spino-thalamic tract ^49,50^. Meta– and mega-analyses of placebo analgesia^51–54^ have found reductions in areas associated with pain processing, including anterior midcingulate (aMCC), medial and ventrolateral thalamus, and anterior and (less consistently) dorsal posterior insula (aIns and dpIns). Reductions in these areas correlate with behavioral placebo analgesia ^53,54^.

However, some evidence suggest that many of these placebo-induced reductions in ‘pain processing’ may be related to affective and decision-making processes rather than nociception, implying a later stage of influence. aMCC and aIns signals are influenced by a wide variety of processes, from emotion to language to motor control ^55–57^. dpIns is the cortical area most selective for nociception ^58–60^, but it also responds to non-somatic, emotional stimuli in some cases ^60,61^. Thus, placebo influences on these regions do not necessarily entail effects on nociception. Stronger tests can be provided by multivariate brain signatures, or neuromarkers ^62^, that can track noxious stimulus intensity and pain with substantially higher sensitivity (larger effect sizes) and specificity than individual regions ^63–65^. In particular, the Neurologic Pain Signature (NPS) ^66^ tracks the intensity of nociceptive input and predicts reported pain with very large effect sizes across cohorts (e.g., d = 1.45 ^67^ or 2.3 ^54^)–and is also largely specific to nociceptive pain: It does not respond to multiple types of non-nociceptive affective stimuli ^65,68–70^. Tests of placebo effects on the NPS in individual studies ^71^ and a recent participant-level meta-analysis across 20 studies (603 participants) ^54^ show little influence of placebo effects. While a significant reduction with placebo was found, its effect size was very small (d = 0.08) compared with the robust behavioral placebo analgesia (d = 0.66), paralleling earlier findings ^72^ showing that neural placebo effects on noxious stimulus-evoked electrophysiological potentials were significant but too small to explain the behavioral analgesic effects of placebo. Small and condition-selective effects on early sensory construction make sense from a functional perspective as well. While early sensory modulation could be adaptive and energy efficient ^22,24^, it could also be dangerous–leading to false perceptions and hallucinations–and could impair learning by reducing prediction errors and error-driven learning in post-sensory processes, preventing the development of epistemically accurate internal models of the world.

Definitive tests of placebo effects on nociceptive and extra-nociceptive pain-related brain processes require highly powered tests of both positive (significant) and null effects (e.g., using Bayes factors ^73^) in large samples, and the assessment of effect sizes in a priori markers. Here, we report such tests on the largest single neuroimaging sample of placebo analgesia to date (N = 392 after exclusions). We tested placebo effects on thermal pain reinforced by a placebo response conditioning procedure ^74^ and transfer to unreinforced effects on mechanical pain (Figure 1 and Methods). Signature-based analyses focused on the NPS and a second neuromarker for higher-level endogenous contributions to pain–the Stimulus Intensity independent Pain Signature (SIIPS ^75^)–originally trained and validated across six studies to predict pain ratings after removing variance associated with stimulus intensity and the NPS. Unlike NPS, SIIPS has been found to mediate the effects of several psychological manipulations on pain, including effects of conditioned auditory cues, visual cues, and perceived control ^75^. In addition, we tested three sets of pre-registered brain regions of interest (ROIs; Supplementary Table 1; for pre-registration see https://osf.io/unh7f and Methods). One set focused on regions most closely associated with nociceptive pain, the second one focused on sub-regions of the SIIPS signature, and the third set focused on prefrontal and striatal regions broadly involved in motivation, value, attention and emotion regulation ^50,51,76–79^.

**Figure 1.**
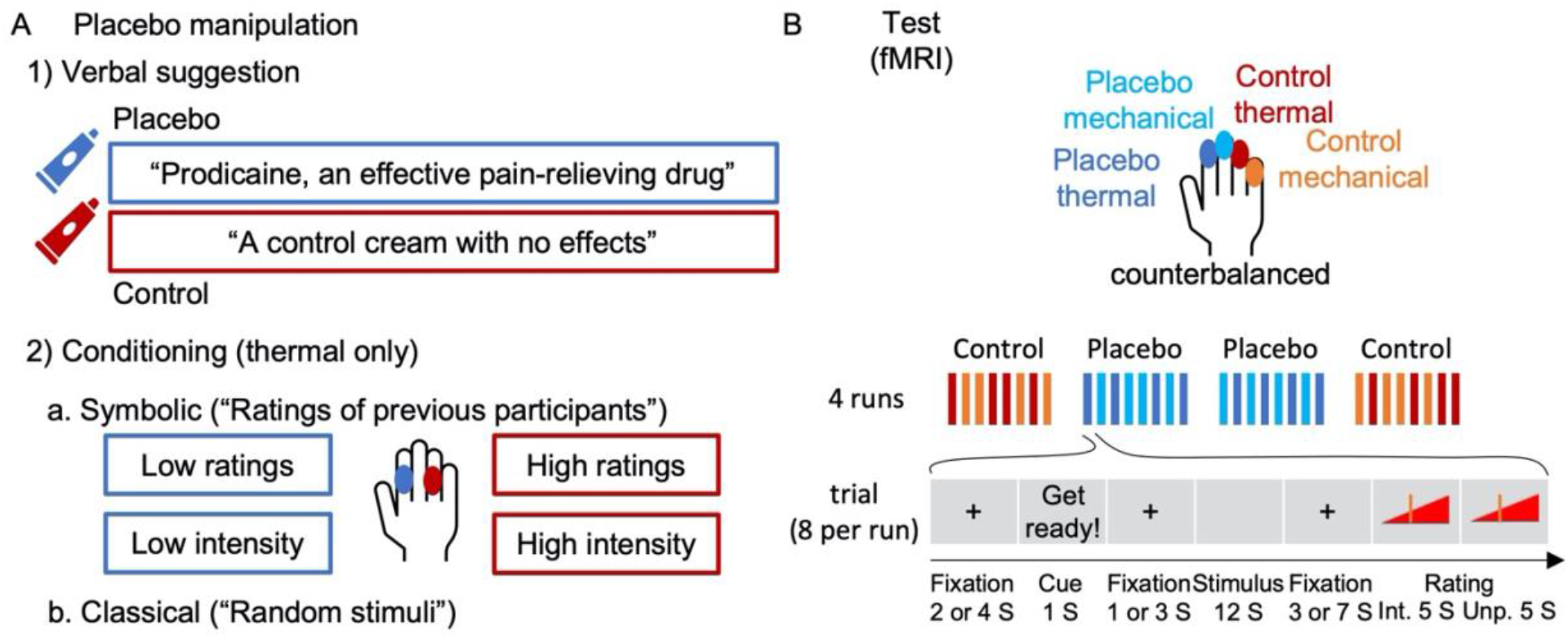
Experimental design. (**A**) First, participants were introduced to the same cream, once presented as “Prodicaine, an effective pain-relieving drug” and once presented as “a control cream with no effects”. Then they went through two conditioning phases: 1) A symbolic conditioning phase, in which “ratings of previous participants” were presented for a series of sham pain trials, with ratings systematically higher for the control compared to the placebo skin site. 2) A classical conditioning phase, in which participants experienced a series of thermal stimuli that they thought were random but in fact were experimentally manipulated to be higher for the control compared to the placebo skin site. (**B**) Then, the test task took place in the MRI scanner. The task consisted of four runs (order: control, placebo, placebo, control), each including eight trials, four with thermal stimulation and four with mechanical (not conditioned) stimulation. Stimuli in the test task were from three intensity levels per modality. In each trial, participants saw a cue, then experienced the stimulus and rated its intensity and unpleasantness. For further details see Methods. Abbreviations: S (seconds); Int. (intensity); Unp. (unpleasantness).

In addition to tests of nociceptive and affective pain neuromarkers and regions, this study addressed an important, unanswered question about transfer (i.e., generalization) of placebo effects. Historical studies have found that placebo effects do not generalize across different types of pain (e.g., labor, postpartum, and experimental, ischemic muscle pain ^80^), and minor variations in context ^81^. Moreover, substantial attempts to identify “placebo responders” across domains have been mostly unsuccessful ^82^. On the other hand, one of the main underlying mechanisms of placebo effects is associative learning, which is known to generalize across stimuli following successful learning ^83–85^. Furthermore, it has been shown that placebo effects transfer across different routes of drug administration ^86^ and over time based on treatment history ^87,88^, and also across domains in some cases (e.g., from pain to negative emotions ^89^ or from pain to motor performance ^90^). In many cases, rather than across domains, placebo effects may transfer between modalities and stimuli within the same domain (e.g., between thermal and mechanical pain, but not from pain to itch ^91^). No or weak placebo effects in the transfer condition would indicate the involvement of specific learning mechanisms in placebo analgesia, whereas strong effects would suggest involvement of inferential and expectancy-related processes ^92–94^.

## Results

### Behavioral placebo analgesia

First, we tested the effect of the placebo manipulation on pain ratings. We included three levels of intensity for each modality (thermal and mechanical pain), selected from pilot testing to be painful and tolerable in a broad population sample (thermal: 46.5, 47 and 47.5 °C; mechanical: 6, 7 and 8 kg/cm^2^). This allowed us to test for stimulus intensity effects, placebo effects, and their interaction on all outcomes, with a mixed-effects model controlling for the familial structure (see Methods). Subjective ratings of stimulus intensity and unpleasantness were highly correlated across trials within participants (*Pearson’s r* was computed for each of 372 participants: across participants, median = 0.956, *M* = 0.896, *sd* = 0.171). Therefore, we focused on intensity ratings, which are typically less responsive than unpleasantness ratings to psychological interventions ^95^. The same pattern of effects was found with the unpleasantness ratings (see Supplementary Information). The ratings were provided on a generalized Labeled Magnitude Scale (gLMS ^96^; 0 = no pain; 0.014 = barely detectable; 0.061 = weak pain; 0.172 = moderate pain; 0.354 = strong pain; 0.533 = very strong pain; 1 = most intense sensation imaginable).

### Thermal pain

As shown in Figure 2, thermal pain ratings increased with stimulus intensity (*M* = 0.129, 0.148, and 0.191 for low, medium, and high intensity; Intensity effect: β = 0.409, *SE* = 0.031, *t*_(416.2)_ = 13.32, *p* < .001, 95% CI = [0.348, 0.469]) and were lower in the Placebo compared to the Control condition (Placebo *M* = 0.129, Control *M* = 0.183; Placebo effect: β = –0.359, *SE* = 0.037, *t*(230.9) = –9.73, *p* < .001, 95% CI = [-0.432, –0.286]). The Intensity x Placebo interaction was not significant (β = –0.080, *SE* = 0.053, *t*_(1282.9)_ = –1.51, *p* = .131, 95% CI = [-0.183, 0.024]). These results were robust to the inclusion of demographic covariates (sex and age, see “Robustness to covariates and demographic effects” in the Supplementary Information). In addition, individual differences in placebo analgesia were significantly correlated with pre-scan ratings of expected Prodicaine efficacy (β = 0.107, *SE* = 0.036, *t*_(339)_ = 3.00, *p* = .003, 95% CI = [0.368, 0.176]), indicating that participants who expected higher efficacy experienced stronger placebo analgesia.

**Figure 2.**
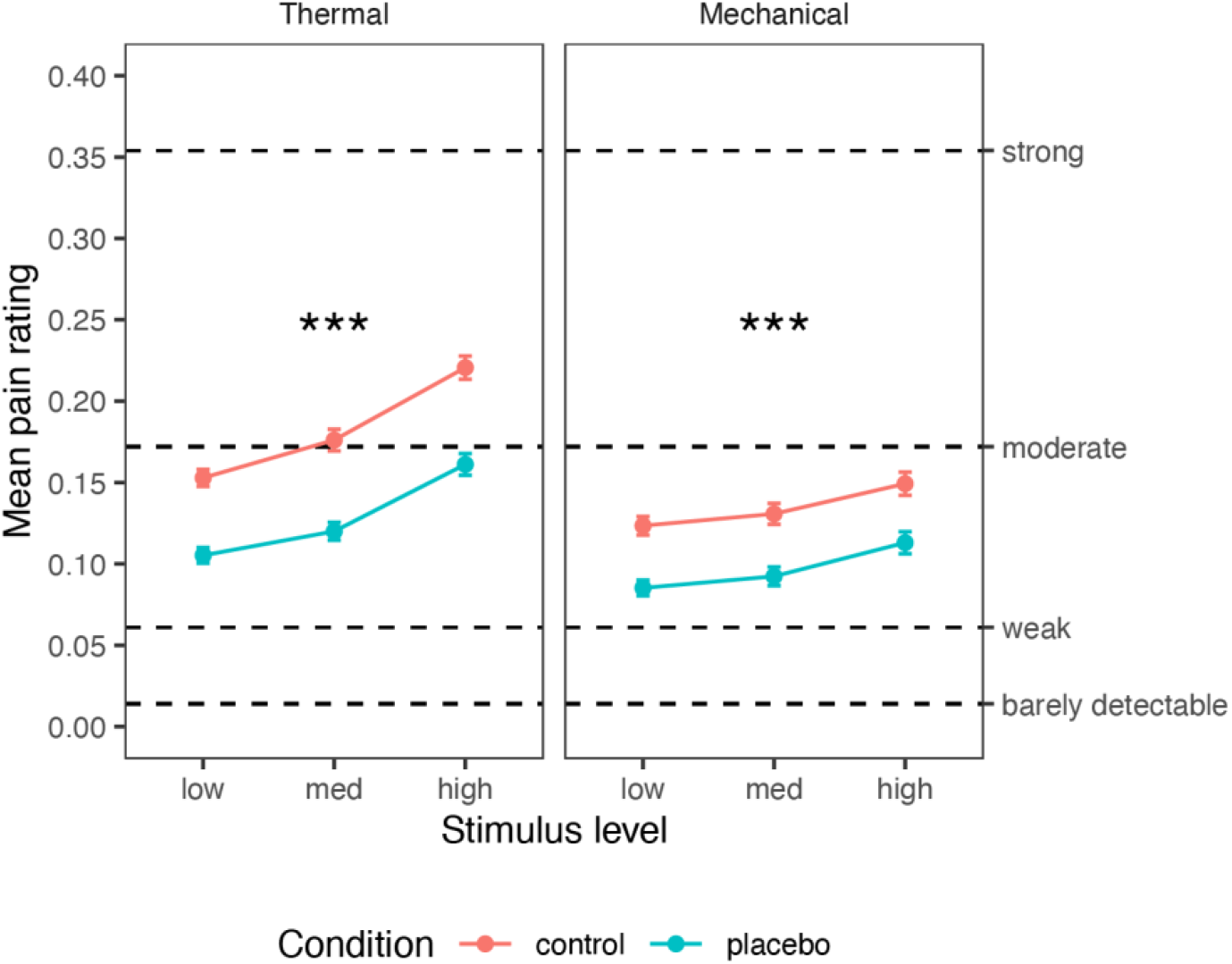
Behavioral results. Mean pain ratings for Control (red) and Placebo (blue) condition, for each combination of modality and stimulus level. Error bars represent within-participant standard error of the mean, based on Morey, 2008 ^97^. Asterisks represent significance of the placebo effect (Placebo vs. Control, uncorrected): * *p* < .05, ** *p* < .01, *** *p* < .001. For additional visualizations see Supplementary Figure 1.

### Mechanical pain

Although the placebo cream (Prodicaine) was only conditioned with the thermal stimuli, its effect transferred from the conditioned thermal modality to the unconditioned mechanical modality. Mechanical pain ratings increased with Stimulus Intensity (*M* = 0.104, 0.112, and 0.131; Intensity effect: β = 0.202, *SE* = 0.032, *t*_(121.7)_ = 6.39, *p* < .001, 95% CI = [0.139, 0.264]) and decreased with Placebo (Placebo *M* = 0.097, Control *M* = 0.134; Placebo effect: β = –0.244, *SE* = 0.039, *t*_(345.1)_ = –6.24, *p* < .001, 95% CI = [-0.321, –0.167]), with no significant interaction (β = 0.013, *SE* = 0.038, *t*_(1089.2)_ = 0.33, *p* = .741, 95% CI = [-0.062, 0.087]). These results were robust to the inclusion of demographic covariates (Supplementary Information, “Robustness to covariates and demographic effects”). Pre-scan expectations of Prodicaine efficacy were not correlated with the placebo effect in the mechanical modality (β = 0.010, *SE* = 0.044, *t*_(163.5)_ = 0.23, *p* = .817, 95% CI = [-0.077, 0.097]).

Furthermore, there was no significant difference in the placebo effect between the thermal and mechanical modalities (β = –0.005, *SE* = 0.051, *t*_(202)_ = –0.09, *p* = .925, 95% CI = [-0.106, 0.096]). Placebo effects were positively correlated across thermal and mechanical modalities, indicating that participants with stronger placebo effects in the thermal condition also showed a stronger placebo effect in the mechanical modality (β = 0.215, *SE* = 0.071, *t*_(151.2)_ = 3.03, *p* = .003, 95% CI = [0.075, 0.355]), providing additional evidence for stable placebo responses across pain types and conditioned vs. transfer modalities.

### Placebo effects on nociceptive processes

#### Placebo effects on the Neurologic Pain Signature (NPS)

The NPS (Figure 3A ^66^) served as a neuromarker for nociceptive pain-related processes based on previous work. As described in the introduction, the NPS is highly sensitive and specific to nociceptive pain.

#### Thermal pain

As shown in Figure 3B, replicating previous findings, the NPS score was positive during thermal stimuli (Cohen’s d = 1.11), and increased with increasing stimulus intensity (β = 0.245, *SE* = 0.041, *t*_(644.5)_ = 6.00, *p* < .001, 95% CI = [0.165, 0.325]). However, the NPS response was not significantly affected by placebo treatment (β = –0.030, *SE* = 0.036, *t*_(225.9)_ = –0.83, *p* = .408, 95% CI = [-0.101, 0.041]), and the Intensity X Placebo interaction was not significant (β = 0.074, *SE* = 0.078, *t*_(1351.5)_ = 0.95, *p* = .342, 95% CI = [-0.079, 0.227]). A Bayes Factor (BF) analysis revealed strong evidence in favor of the null hypothesis of no placebo effect (BF = 0.044, 23:1 odds in favor of the null, proportional error estimate = 9.80%). Since there is some disagreement on certain analytical choices in the context of Bayes Factor analysis ^98^, we further explored the robustness of these results to different analytical choices, particularly with regard to the width of the prior distribution and inclusion of interaction terms. Such variations did not change the conclusion, with all models indicating strong to extreme evidence in favor of the null (BF range 0.036-0.069; Supplementary Table 2). Furthermore, pre-scan expectations of Prodicaine efficacy were not correlated with the placebo effect on the NPS score (β = 0.033, *SE* = 0.034, *t*_(230.8)_ = 0.99, *p* = .323, 95% CI = [-0.033, 0.100]).

**Figure 3.**
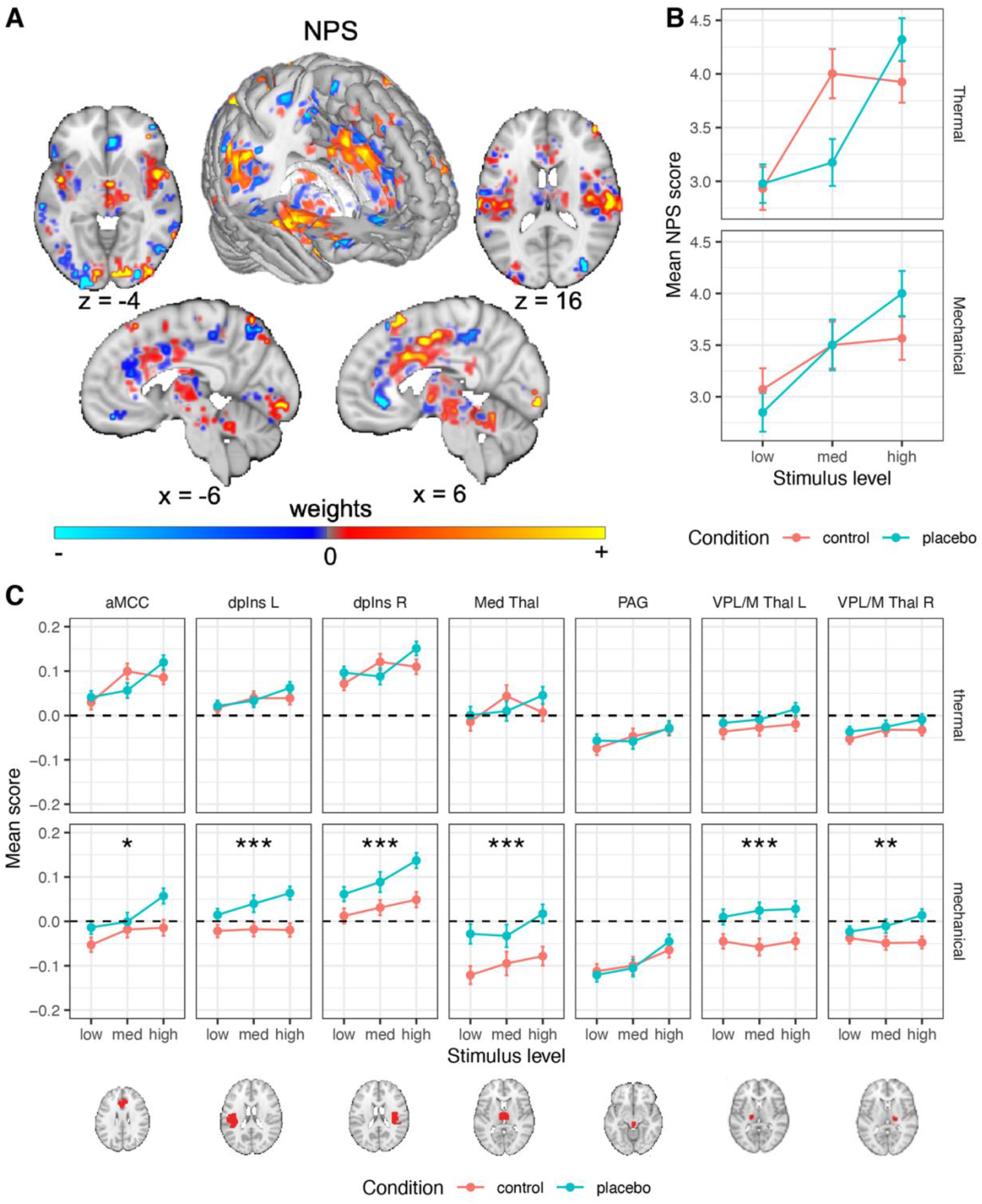
Neural results: a priori nociceptive neuromarker and ROIs. (**A**) The NPS, an fMRI measure optimized to predict pain intensity ^66^. Dominant model parameters are highlighted by transparency scaling. (**B**) The mean NPS scores (based on dot product) across participants are presented, in the Control (red) and Placebo (blue) condition, for each combination of modality and stimulus level. For additional visualizations see Supplementary Figure 2. (**C**) The mean signal across all voxels of each a priori nociceptive ROI across participants are presented, in the Control (red) and Placebo (blue) condition, for each combination of modality and stimulus level. Error bars (in panels B and D) represent within-participant standard error of the mean, based on Morey, 2008 ^97^. Asterisks represent significance of the placebo effect (placebo vs. control, uncorrected): * *p* < .05, ** *p* < .01, *** *p* < .001. For additional visualizations see Supplementary Figure 3. Regions abbreviation: L (left); R (right); NPS (Neurologic Pain Signature); aMCC (anterior midcingulate cortex); dpIns (dorsal posterior insula); Med Thal (medial thalamus); VPL/M Thal (ventral posterior thalamus); PAG (periaqueductal gray).

#### Mechanical pain

The NPS scores were positive during mechanical pain (Cohen’s d = 1.02), and increased with increased stimulus intensity (β = 0.164, *SE* = 0.041, *t*_(804.1)_ = 4.02, *p* < .001, 95% CI = [0.084, 0.244]), but were not affected by placebo (β = 0.008, *SE* = 0.037, *t*_(308.7)_ = 0.23, *p* = .822, 95% CI = [-0.064, 0.081]; Figure 3B), and the Intensity X Placebo interaction was not significant (β = 0.132, *SE* = 0.079, *t*_(1529)_ = 1.67, *p* = .095, 95% CI = [-0.023, 0.287]). Bayes Factor analysis provided strong evidence against the presence of a placebo effect on the NPS response (BF = 0.036, 28:1 odds in favor of the null, proportional error estimate = 11.7%), and this result was robust to analytical variations with strong to extreme evidence across models (BF range 0.001 – 0.049; Supplementary Table 2). As in the thermal modality, pre-scan expectations of Prodicaine efficacy were not correlated with the placebo effect on the NPS score in mechanical trials (β = 0.012, *SE* = 0.033, *t*_(351.8)_ = 0.38, *p* = .703, 95% CI = [-0.051, 0.076]).

Together, these analyses indicated that placebo does not affect fMRI activity in the most widely validated neuromarker to date for nociceptive pain. These results were robust to the inclusion of demographic covariates (Supplementary Information, “Robustness to covariates and demographic effects”).

### A priori regions of interest

The NPS is only one measure and does not capture all aspects of pain processing. Thus, to further assess effects on regions associated with nociception, we tested effects of placebo, stimulus intensity, and their interaction in a set of seven a priori brain regions associated with nociception (six of which were pre-registered, see methods; for full statistics see Table 1). Because false negatives and false positives here were equally important, we report results based on *p* < .05 without correcting for multiple comparisons across regions. This is because of the nature of the current study, focusing largely on testing pre-registered neural signatures and regions identified in previous literature regions with a substantially larger sample size. Nevertheless, to be slightly more conservative, we note when a result does not survive Bonferroni correction within its set of regions (e.g., within the set of seven nociceptive regions).

#### Thermal pain

Of these regions, the heat-evoked response increased with stimulus intensity in the anterior midcingulate cortex (aMCC), right (contralateral) dpIns, left (ipsilateral) dpIns and PAG (the effects in the PAG and left dpIns do not survive Bonferroni correction), but not in the right and left ventral posterior (VPL/M) or medial thalamus (Table 1). The placebo treatment did not significantly modulate the response to painful heat in any of these regions (all *p*s > 0.11, Table 1; Figure 3C). Bayes Factor analysis revealed strong evidence in favor of the null hypothesis (no effect of placebo) in all these a priori brain nociceptive regions, except the left VPL/M thalamus, which showed moderate evidence for the null (Supplementary Table 2). Again, these results were robust to the inclusion of interaction terms (leading to stronger evidence in favor of the null model) and width of the prior distribution (leading to slightly weaker evidence for some regions, yet still moderate to very strong evidence in favor of the null for all regions; all BFs < 0.143). In addition, Intensity x Placebo interactions were non-significant in all regions. These results provide strong evidence that placebo treatment does not modulate response to thermal stimuli in these a priori nociceptive brain regions.

#### Mechanical pain

Pressure-evoked responses increased with stimulus intensity in the right (contralateral) dpIns, PAG and aMCC, but not in the left dpIns, right and left VPL/M thalamus, and medial thalamus (Table 1). Surprisingly, in the mechanical pain transfer condition, placebo treatment increased activity in all these a priori nociception-related regions compared with the control condition, except for the PAG (*p* = .012 or lower across individual regions; Table 1 and Figure 3C). Bayes Factor analysis indicated extreme evidence in favor of the alternative hypothesis (increased activity during placebo compared to control) in the left and right dpIns, left VPL/M thalamus and medial thalamus, strong evidence in the right VPL/M thalamus, and moderate evidence in the aMCC (these results were robust to the width of the prior distribution, but not to the inclusion of an interaction term; Supplementary Table 2). In the PAG, the Bayes Factor analysis provided strong evidence in favor of the null hypothesis (no placebo effect). Intensity x Placebo interactions were all non-significant.

**Table 1.**
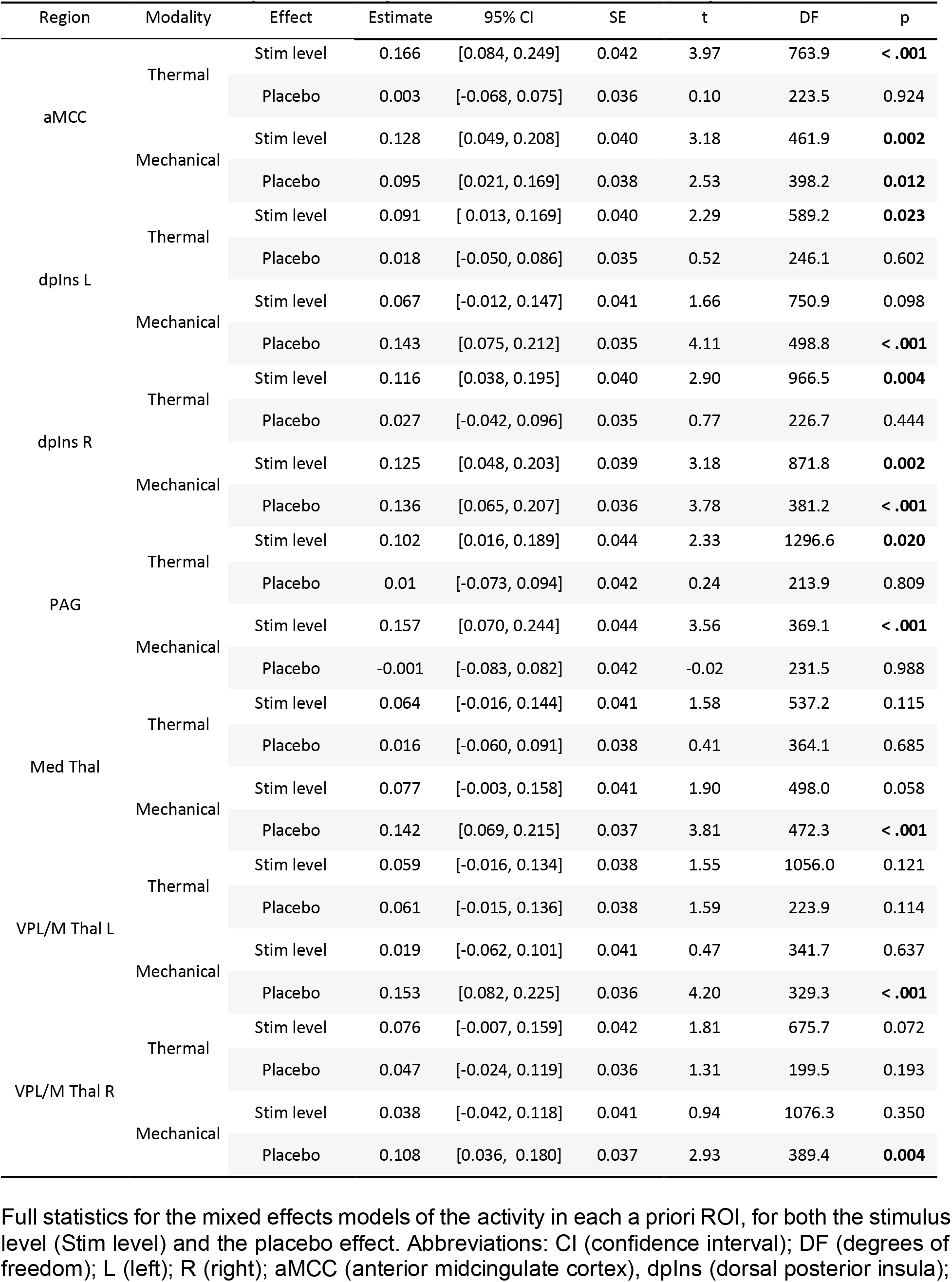
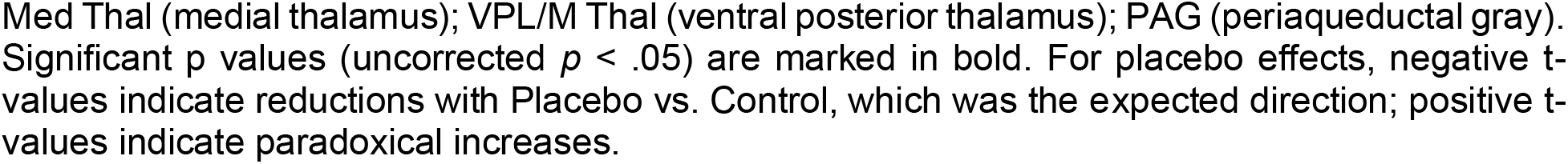
Statistics for a priori nociceptive ROIs: stimulus level and placebo effects.

| Region | Modality | Effect | Estimate | 95% CI | SE | t | DF | p |
| --- | --- | --- | --- | --- | --- | --- | --- | --- |
| aMCC | Thermal | Stim level | 0.166 | [0.084, 0.249] | 0.042 | 3.97 | 763.9 | **< .001** |
|  |  | Placebo | 0.003 | [-0.068, 0.075] | 0.036 | 0.10 | 223.5 | 0.924 |
|  | Mechanical | Stim level | 0.128 | [0.049, 0.208] | 0.040 | 3.18 | 461.9 | **0.002** |
|  |  | Placebo | 0.095 | [0.021, 0.169] | 0.038 | 2.53 | 398.2 | **0.012** |
| dplns L | Thermal | Stim level | 0.091 | [0.013, 0.169] | 0.040 | 2.29 | 589.2 | **0.023** |
|  |  | Placebo | 0.018 | [-0.050, 0.086] | 0.035 | 0.52 | 246.1 | 0.602 |
|  | Mechanical | Stim level | 0.067 | [-0.012, 0.147] | 0.041 | 1.66 | 750.9 | 0.098 |
|  |  | Placebo | 0.143 | [0.075, 0.212] | 0.035 | 4.11 | 498.8 | **< .001** |
| dplns R | Thermal | Stim level | 0.116 | [0.038, 0.195] | 0.040 | 2.90 | 966.5 | **0.004** |
|  |  | Placebo | 0.027 | [-0.042, 0.096] | 0.035 | 0.77 | 226.7 | 0.444 |
|  | Mechanical | Stim level | 0.125 | [0.048, 0.203] | 0.039 | 3.18 | 871.8 | **0.002** |
|  |  | Placebo | 0.136 | [0.065, 0.207] | 0.036 | 3.78 | 381.2 | **< .001** |
| PAG | Thermal | Stim level | 0.102 | [0.016, 0.189] | 0.044 | 2.33 | 1296.6 | **0.020** |
|  |  | Placebo | 0.01 | [-0.073, 0.094] | 0.042 | 0.24 | 213.9 | 0.809 |
|  | Mechanical | Stim level | 0.157 | [0.070, 0.244] | 0.044 | 3.56 | 369.1 | **< .001** |
|  |  | Placebo | -0.001 | [-0.083, 0.082] | 0.042 | -0.02 | 231.5 | 0.988 |
| Med Thal | Thermal | Stim level | 0.064 | [-0.016, 0.144] | 0.041 | 1.58 | 537.2 | 0.115 |
|  |  | Placebo | 0.016 | [-0.060, 0.091] | 0.038 | 0.41 | 364.1 | 0.685 |
|  | Mechanical | Stim level | 0.077 | [-0.003, 0.158] | 0.041 | 1.90 | 498.0 | 0.058 |
|  |  | Placebo | 0.142 | [0.069, 0.215] | 0.037 | 3.81 | 472.3 | **< .001** |
| VPL/M Thal L | Thermal | Stim level | 0.059 | [-0.016, 0.134] | 0.038 | 1.55 | 1056.0 | 0.121 |
|  |  | Placebo | 0.061 | [-0.015, 0.136] | 0.038 | 1.59 | 223.9 | 0.114 |
|  | Mechanical | Stim level | 0.019 | [-0.062, 0.101] | 0.041 | 0.47 | 341.7 | 0.637 |
|  |  | Placebo | 0.153 | [0.082, 0.225] | 0.036 | 4.20 | 329.3 | **< .001** |
| VPL/M Thal R | Thermal | Stim level | 0.076 | [-0.007, 0.159] | 0.042 | 1.81 | 675.7 | 0.072 |
|  |  | Placebo | 0.047 | [-0.024, 0.119] | 0.036 | 1.31 | 199.5 | 0.193 |
|  | Mechanical | Stim level | 0.038 | [-0.042, 0.118] | 0.041 | 0.94 | 1076.3 | 0.350 |
|  |  | Placebo | 0.108 | [0.036, 0.180] | 0.037 | 2.93 | 389.4 | **0.004** |

Full statistics for the mixed effects models of the activity in each a priori ROI, for both the stimulus level (Stim level) and the placebo effect. Abbreviations: CI (confidence interval); DF (degrees of freedom); L (left); R (right); aMCC (anterior midcingulate cortex), dplns (dorsal posterior insula);
Med Thal (medial thalamus); VPL/M Thal (ventral posterior thalamus); PAG (periaqueductal gray). Significant $p$ values (uncorrected $p < .05$ ) are marked in bold. For placebo effects, negative $t$ -values indicate reductions with Placebo vs. Control, which was the expected direction; positive $t$ -values indicate paradoxical increases.

Since there were several surprising local increased activations for Placebo compared to Control during mechanical pain stimuli specifically, we performed exploratory analysis to test whether these effects can be accounted for by carry-over effects from the anticipation period. Anticipation responses did not account for these effects, as anticipatory brain activity in these regions did not significantly differ between Placebo and Control runs for the mechanical trials, except for the aMCC where anticipatory activity in mechanical trials was significantly lower in Placebo compared to Control (β = –0.098, *SE* = 0.049, *t*(287.4) = –2.00, *p* = .047, 95% CI = [-0.195, –0.001]). A full analysis of the anticipation data is beyond the scope of the current paper and will be reported in future ones.

Taken together, these findings provide definitive evidence that placebo analgesia does not reduce activity in regions associated with nociceptive pain on average in this study.

### Placebo effects on higher-level processes

Pain perception is more than nociceptive processing, and includes higher-level cognitive and affective processes. Thus, we also tested the effect of placebo on such higher-level pain processing, including a priori neuromarker and regions of interest from previous studies.

#### A priori neuromarker

First, we used SIIPS ^75^, a neuromarker trained to predict pain independent of stimulus intensity and the NPS. SIIPS was designed to capture higher-level, endogenous brain influences on pain construction, beyond the nociceptive effects captured by the NPS. This signature was trained on four different datasets (*N* = 137 participants overall) and tested on two independent datasets (*N* = 46), explaining trial-level variance in pain ratings beyond the variance explained by the input intensity and NPS, and mediating the effects of three psychological manipulations on pain (two eliciting expectancy effects and one manipulating perceived control). SIIPS includes specific activity patterns in key placebo-linked regions, including vmPFC, dlPFC, hippocampus, and nucleus accumbens (NAc).

##### Thermal pain

As predicted, the SIIPS stimulus-evoked response was significantly reduced by the placebo treatment (β = –0.129, *SE* = 0.034, *t*_(417.4)_ = –3.83, *p* < .001, 95% CI = [-0.195, –0.063]), indicating that the endogenous processes captured by SIIPS are modulated by placebo effects (Figure 4A). Bayes Factor analysis provided extreme evidence for this conclusion (*BF* = 93.92, *proportional error* = 13.9%; extreme evidence across variations beside inclusion of an interaction term; Supplementary Table 3).

Though SIIPS was trained to explain variance in pain reports after controlling for stimulus intensity and nociceptive processes, it has been found to respond more to stronger stimuli in previous studies ^75^. This can occur if variations in some sub-regions of SIIPS that are considered nociceptive (e.g., parts of the insula and cingulate cortex) contribute to pain beyond intensity encoding. Indeed, the SIIPS stimulus-evoked response was also stronger for more intense stimuli in the current study (β = 0.181, *SE* = 0.038, *t*(1408.3) = 4.77, *p* < .001, 95% CI = [0.106, 0.255]). Intensity x Placebo interaction was non-significant (β = –0.055, *SE* = 0.075, *t*_(1559.9)_ = –0.73, *p* = .467, 95% CI = [-0.203, 0.093]). Pre-scan expectations of Prodicaine efficacy were not correlated with the placebo effect on the SIIPS score (β = 0.042, *SE* = 0.031, *t*_(251.4)_ = 1.35, *p* = .178, 95% CI = [-0.019, 0.103]).

##### Mechanical pain

The effect of placebo on the SIIPS stimulus-evoked response transferred to the unconditioned mechanical pain modality, with lower SIIPS response in the Placebo compared to the Control treatment (β = –0.112, *SE* = 0.037, *t*_(286.1)_ = –3.00, *p* = .003, 95% CI = [-0.185, – 0.039]; Figure 4A). This effect was not significantly different from the placebo effect in the thermal modality (β = 0.0002, *SE* = 0.045, *t*_(213.5)_ = 0.005, *p* = .996, 95% CI = [-0.088, 0.088]). As in the thermal modality, the SIIPS response was also stronger for more intense mechanical stimuli (β = 0.188, *SE* = 0.039, *t*_(233)_ = 4.75, *p* < .001, 95% CI = [0.110, 0.265]). The Intensity x Placebo interaction was non-significant (β = 0.090, *SE* = 0.071, *t*_(1274.2)_ = 1.27, *p* = .205, 95% CI = [-0.049, –0.229]). The SIIPS results were robust to the inclusion of demographic covariates (Supplementary Information, “Robustness to covariates and demographic effects”). Like in the thermal modality, pre-scan expectations of Prodicaine efficacy were not correlated with the placebo effect on the SIIPS score in mechanical trials (β = –0.021, *SE* = 0.035, *t*_(299.3)_ = –0.61, *p* = .539, 95% CI = [-0.090, 0.047]).

**Figure 4.**
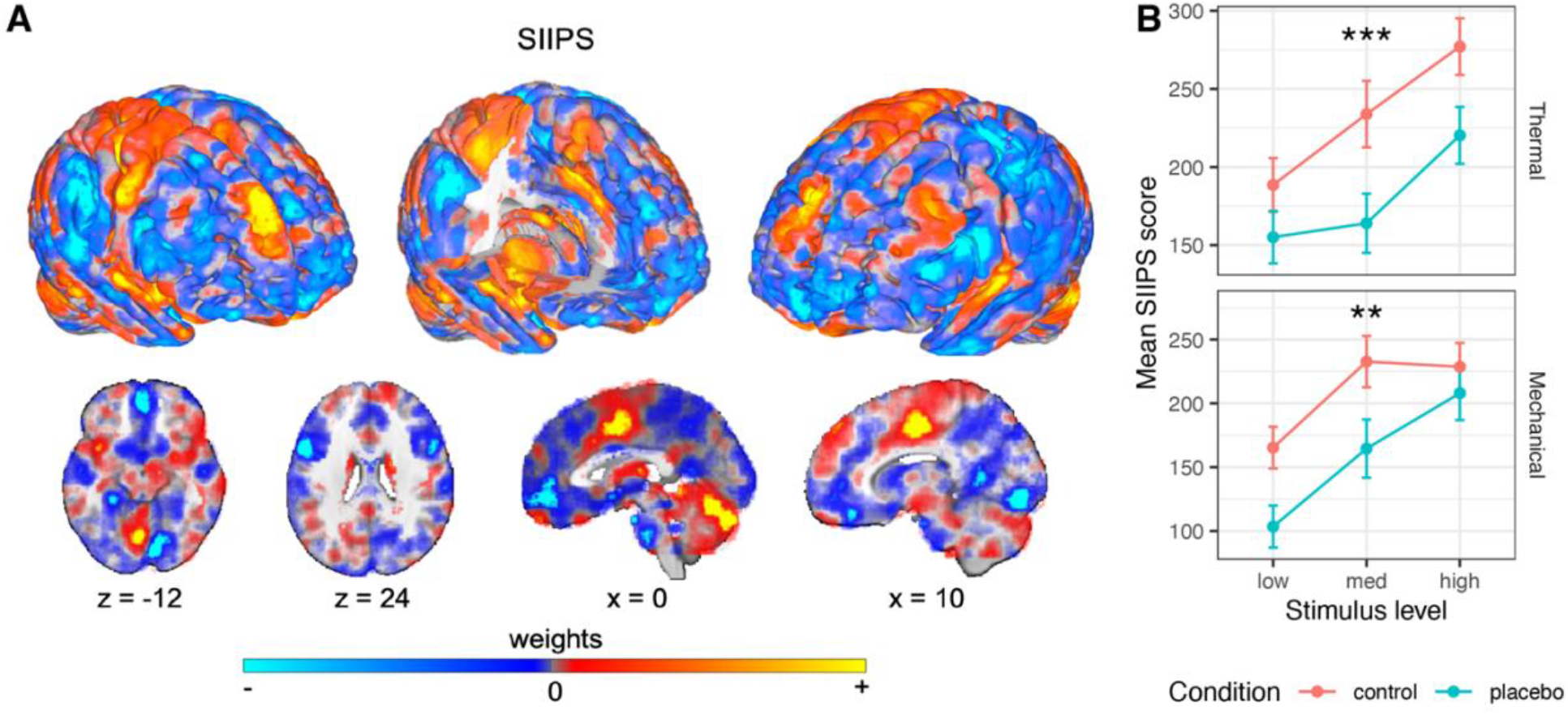
Neural results: a priori higher-level pain processing neuromarker. (**A**) The SIIPS signature, fMRI measure optimized to predict pain beyond nociception ^75^. Dominant model parameters are highlighted by transparency scaling. (**B**) The mean SIIPS scores (based on dot product) across participants are presented, in the Control (red) and Placebo (blue) condition, for each combination of modality and stimulus level. Error bars represent within-participant standard error of the mean, based on Morey, 2008 ^97^. For additional visualizations see Supplementary Figure 4. Asterisks represent significance of the placebo effect (uncorrected): * *p* < .05, ** *p* < .01, *** *p* < .001. For visualizations of SIIPS subregions see Supplementary Figures 6 and 7.

##### Subregions

SIIPS is a multivariate signature that includes positive and negative weights across the brain. A subset of regions in the brain have robust pattern weights (i.e., they made consistent contributions to prediction across participants and studies in the original paper ^75^). These subregions can be divided into three types: (1) Regions with positive weights (i.e., increased activity is associated with more pain) and which are established as targets of nociceptive inputs. These regions include the insula, thalamus and cingulate cortex. They overlap with some of the gross anatomical regions included in the NPS, however they reflect distinct local regions, and their weights in SIIPS are not correlated with their weights in the NPS. Thus, they likely encode pain beyond the nociceptive processes captured by NPS. (2) Regions with positive weights and which are not known to be nociceptive, including the dmPFC and caudate. These regions likely encode pain processing beyond nociception. (3) Regions with negative weights (i.e., increased activity in these regions was associated with less pain), including areas in the NAc and the vmPFC. These regions are thought to encode cognitive and affective processes related to pain regulation and other pain-opposing processes, and are thought to play an important role in chronic _pain_ 99–101.

We focused on eight specific subregions of SIIPS that showed non-significant responses to noxious stimulus intensity in the SIIPS’ training studies and are thus considered non-nociceptive contributors to the pain experience. These subregions include two regions with positive weights (dmPFC and right middle temporal gyrus, rMTG) and six regions with negative weights (right lingual gyrus, left superior temporal gyrus [lSTG], left NAc, right temporal pole, left inferior temporal gyrus [lITG] and middle precentral gyrus). In addition to these eight pre-registered subregions of SIIPS, we also tested for effects in seven subregions that were not pre-registered but are of interest based on recent work, including the vmPFC, right nucleus accumbens, left and right dlPFC, right secondary somatosensory cortex, right sensorimotor cortex and left precuneus (for the results in these regions see Supplementary Information, “Additional, non-pre-registered SIIPS subregions”).

For full statistics in all eight pre-registered subregions for both modalities, see Table 2. We report local pattern response in each region, which is the dot product of the activity in each voxel and its corresponding SIIPS pattern weights. The interpretation is somewhat different from the standard average activity reported for ROIs. In regions with positive weights, more positive pattern responses indicate greater activation and increased pain. In subregions with negative weights, positive pattern responses indicate *deactivation* and predict *increased* pain (activity in these regions is also scaled based on the weights across the different voxels). Thus, more positive responses indicated more pain-related activity in all regions, and we expected decreased pattern responses during Placebo compared to Control treatment (negative t-values) in all subregions. In “SIIPS-Pos” regions, decreased pattern response corresponds to decreased overall activity with placebo. In “SIIPS-Neg” regions (e.g., NAc), decreased pattern response corresponds to increased overall activity with placebo.

dmPFC and rMTG (the regions with positive weights) showed no placebo effects in either thermal or mechanical pain (Supplementary Figure 5). The only region with significant placebo-related decreases in local SIIPS pattern response during thermal pain was lSTG (however, this result did not survive Bonferroni correction). On mechanical pain trials, we found significant placeb o-induced decreases in the right lingual gyrus, lSTG, left NAc, middle precentral gyrus and right temporal pole, but not lITG (and the effect in the right temporal pole did not survive Bonferroni correction). Unexpectedly, there was a significant negative effect of stimulus level in the rMTG in the thermal modality, and a significant positive effect of stimulus level in the dmPFC in the mechanical modality, but both effects did not survive Bonferroni correction.

**Table 2.** Statistics for a priori subregions of SIIPS: stimulus level and placebo effects.

| Region | Modality | Effect | Estimate | 95% CI | SE | t | DF | p |
| --- | --- | --- | --- | --- | --- | --- | --- | --- |
| SIIPS Pos - dmPFC | Thermal | Stim level | -0.028 | [-0.106, 0.050] | 0.040 | -0.70 | 998.6 | 0.484 |
|  |  | Placebo | 0.035 | [-0.035, 0.105] | 0.036 | 0.99 | 228.5 | 0.324 |
|  | Mechanical | Stim level | 0.089 | [0.009, 0.168] | 0.041 | 2.19 | 1194.3 | **0.029** |
|  |  | Placebo | 0.078 | [-0.003, 0.158] | 0.041 | 1.91 | 230.9 | 0.058 |
| SIIPS Pos - MTG R | Thermal | Stim level | -0.108 | [-0.197,-0.019] | 0.045 | -2.39 | 773.9 | **0.017** |
|  |  | Placebo | 0.074 | [-0.003, 0.152] | 0.039 | 1.89 | 434.6 | 0.060 |
|  | Mechanical | Stim level | -0.046 | [-0.132, 0.040] | 0.044 | -1.06 | 1389.9 | 0.291 |
|  |  | Placebo | 0.065 | [-0.012, 0.142] | 0.039 | 1.66 | 325.2 | 0.098 |
| SIIPS Neg - NAc L | Thermal | Stim level | 0.023 | [-0.065, 0.112] | 0.045 | 0.52 | 810.0 | 0.604 |
|  |  | Placebo | 0.012 | [-0.059, 0.083] | 0.036 | 0.34 | 851.7 | 0.735 |
|  | Mechanical | Stim level | -0.089 | [-0.179, 0.001] | 0.046 | -1.95 | 806.9 | 0.052 |
|  |  | Placebo | -0.111 | [-0.187,-0.034] | 0.039 | -2.85 | 466.6 | **0.005** |
| SIIPS Neg - LG R | Thermal | Stim level | 0.015 | [-0.065, 0.094] | 0.041 | 0.36 | 528.3 | 0.720 |
|  |  | Placebo | -0.073 | [-0.148, 0.002] | 0.038 | -1.93 | 211.4 | 0.055 |
|  | Mechanical | Stim level | -0.051 | [-0.131, 0.029] | 0.041 | -1.25 | 664.7 | 0.214 |
|  |  | Placebo | -0.165 | [-0.238,-0.093] | 0.037 | -4.47 | 422.3 | **< .001** |
| SIIPS Neg - STG L | Thermal | Stim level | 0.012 | [-0.071, 0.094] | 0.042 | 0.28 | 690.3 | 0.782 |
|  |  | Placebo | -0.069 | [-0.135,-0.002] | 0.034 | -2.03 | 1145.0 | **0.043** |
|  | Mechanical | Stim level | -0.031 | [-0.113, 0.050] | 0.042 | -0.75 | 1213.8 | 0.453 |
|  |  | Placebo | -0.161 | [-0.231,-0.092] | 0.035 | -4.56 | 749.2 | **< .001** |
| SIIPS Neg - TP R | Thermal | Stim level | -0.047 | [-0.133, 0.039] | 0.044 | -1.07 | 1558.6 | 0.286 |
|  |  | Placebo | -0.064 | [-0.138, 0.009] | 0.037 | -1.72 | 239.6 | 0.087 |
|  | Mechanical | Stim level | -0.061 | [-0.154, 0.033] | 0.048 | -1.27 | 341.9 | 0.203 |
|  |  | Placebo | -0.104 | [-0.186,-0.021] | 0.042 | -2.48 | 226.5 | **0.014** |
| SIIPS Neg - ITG L | Thermal | Stim level | -0.069 | [-0.152, 0.015] | 0.042 | -1.62 | 680.2 | 0.106 |
|  |  | Placebo | -0.066 | [-0.136, 0.005] | 0.036 | -1.83 | 505.8 | 0.069 |
|  | Mechanical | Stim level | 0.009 | [-0.079, 0.096] | 0.045 | 0.20 | 1572.1 | 0.846 |
|  |  | Placebo | -0.026 | [-0.105, 0.052] | 0.040 | -0.66 | 214.9 | 0.511 |
| SIIPS Neg -<br>Mid Precen | Thermal | Stim level | 0.034 | [-0.045, 0.113] | 0.040 | 0.84 | 848.9 | 0.401 |
|  |  | Placebo | -0.029 | [-0.095, 0.036] | 0.033 | -0.88 | 323.6 | 0.381 |
|  | Mechanical | Stim level | 0.016 | [-0.063, 0.096] | 0.040 | 0.40 | 516.6 | 0.688 |
|  |  | Placebo | -0.156 | [-0.228,-0.085] | 0.036 | -4.30 | 386.6 | **< .001** |

Full statistics for the mixed effects models of the activity in each a priori subregion of SIIPS, for both the stimulus level (Stim level) and the placebo effect. Abbreviations: CI (confidence interval); DF (degrees of freedom); Pos (positive); Neg (negative); L (left); R (right); dmPFC (dorsomedial prefrontal cortex); MTG (middle temporal gyrus); NAc (nucleus accumbens); LG (lingual gyrus); STG (superior temporal gyrus); TP (temporal pole); ITG (inferior temporal gyrus); mid precen (middle precentral gyrus). Significant p values (uncorrected $p < .05$ ) are marked in bold. For placebo effects, negative coefficients indicate reductions with Placebo vs. Control, which was the expected direction of effects.

#### A priori regions of interest

We tested ten a priori brain regions where increased response to noxious stimuli in placebo compared to control was found in previous studies. These regions are thus suggested to reflect higher-level processes that modulate the placebo effect, such as motivation, emotion regulation, and decision-making. These regions included the right dlPFC, two different areas of the left dlPFC (one more anterior than the other), two areas of the anterior orbitofrontal cortex (OFC; one more inferior and one more superior), lateral OFC, right and left middle lateral OFC, and right and left NAc (here, a simple region average, in contrast to the local pattern responses reported for SIIPS subregions). For full statistics for each individual region, see Table 3.

**Figure 5.**
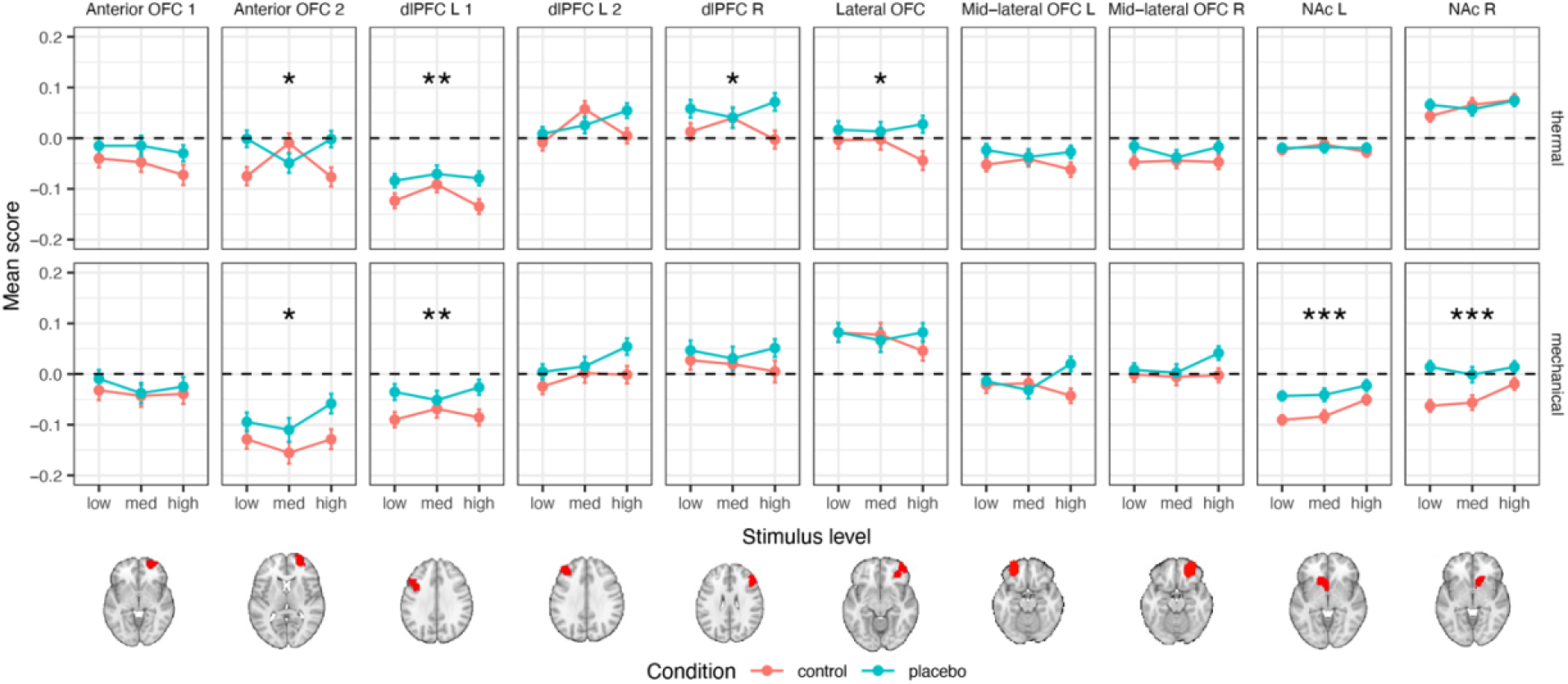
Neural results: a priori higher-level pain processing ROIs. The mean signal across all voxels of each a priori ROI across participants are presented, in the Control (red) and Placebo (blue) condition, for each combination of modality and stimulus level. Error bars represent within-participant standard error of the mean, based on Morey, 2008 ^97^. Asterisks represent significance of the placebo effect (uncorrected): * *p* < .05, ** *p* < .01, *** *p* < .001. For additional visualizations see Supplementary Figure 7. Abbreviations: L (left); R (right); dlPFC (dorsolateral prefrontal cortex); OFC (orbitofrontal cortex); NAc (nucleus accumbens).

##### Thermal pain

We found significant placebo-induced increases in the response to thermal stimuli in the more posterior of the two left dlPFC regions, the right dlPFC, the more superior of the two anterior OFC regions, and the lateral OFC (but only the left dlPFC survives Bonferroni correction; Table 3 and Figure 5). There were no significant placebo effects in the right and left NAc, and in the second (more anterior) a priori area of the left dlPFC, the second (more inferior) a priori area of the anterior OFC, and the right and left mid-lateral OFC.

##### Mechanical pain

During mechanical pain trials, we found significant placebo-induced increases in the more posterior region of the left dlPFC and the more superior region of the anterior OFC (as in thermal pain, though the latter result does not survive Bonferroni correction) and in the left and right NAc (Table 3 and Figure 5). No significant placebo effects were found in the other a priori regions.

**Table 3.**
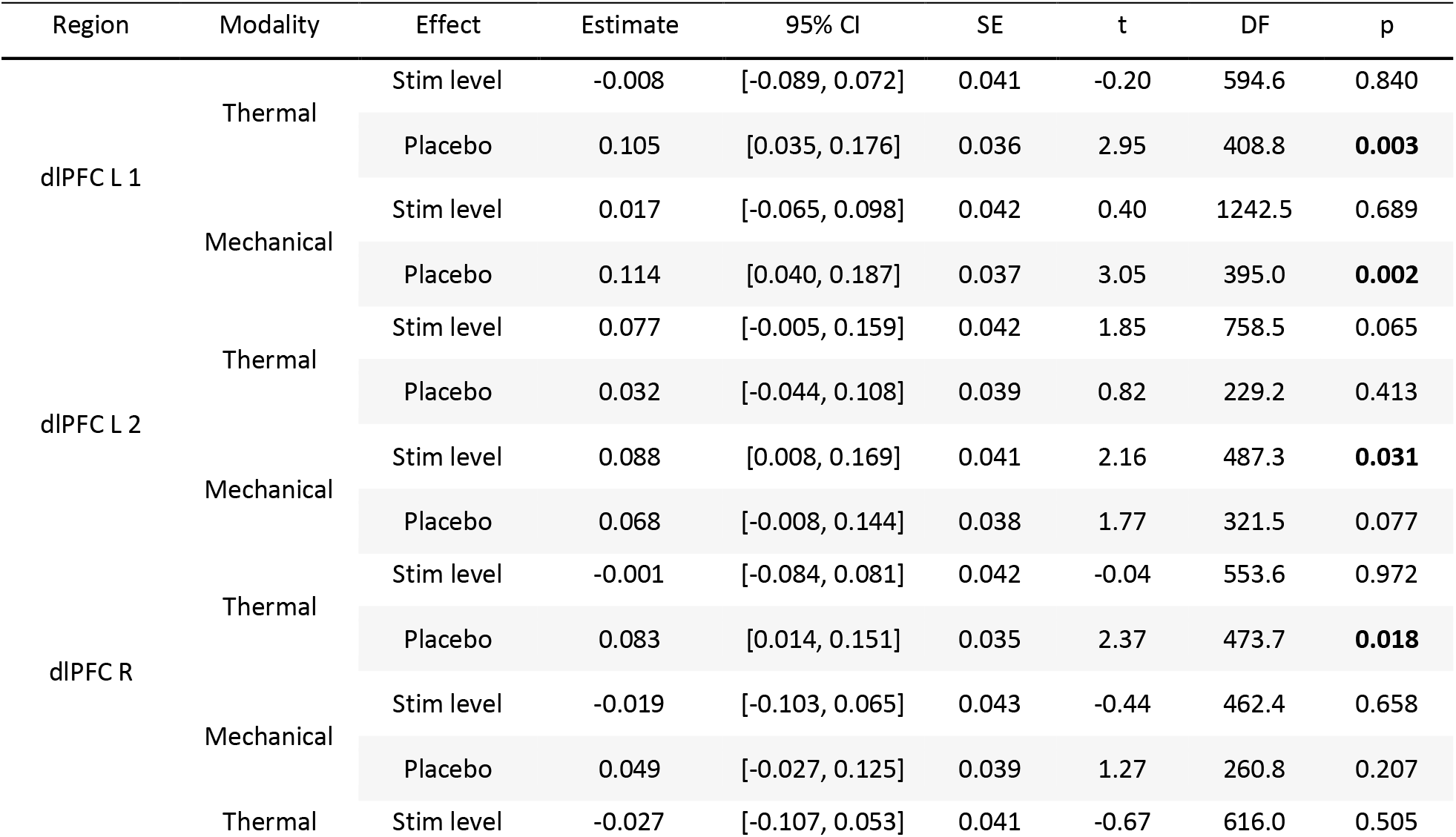
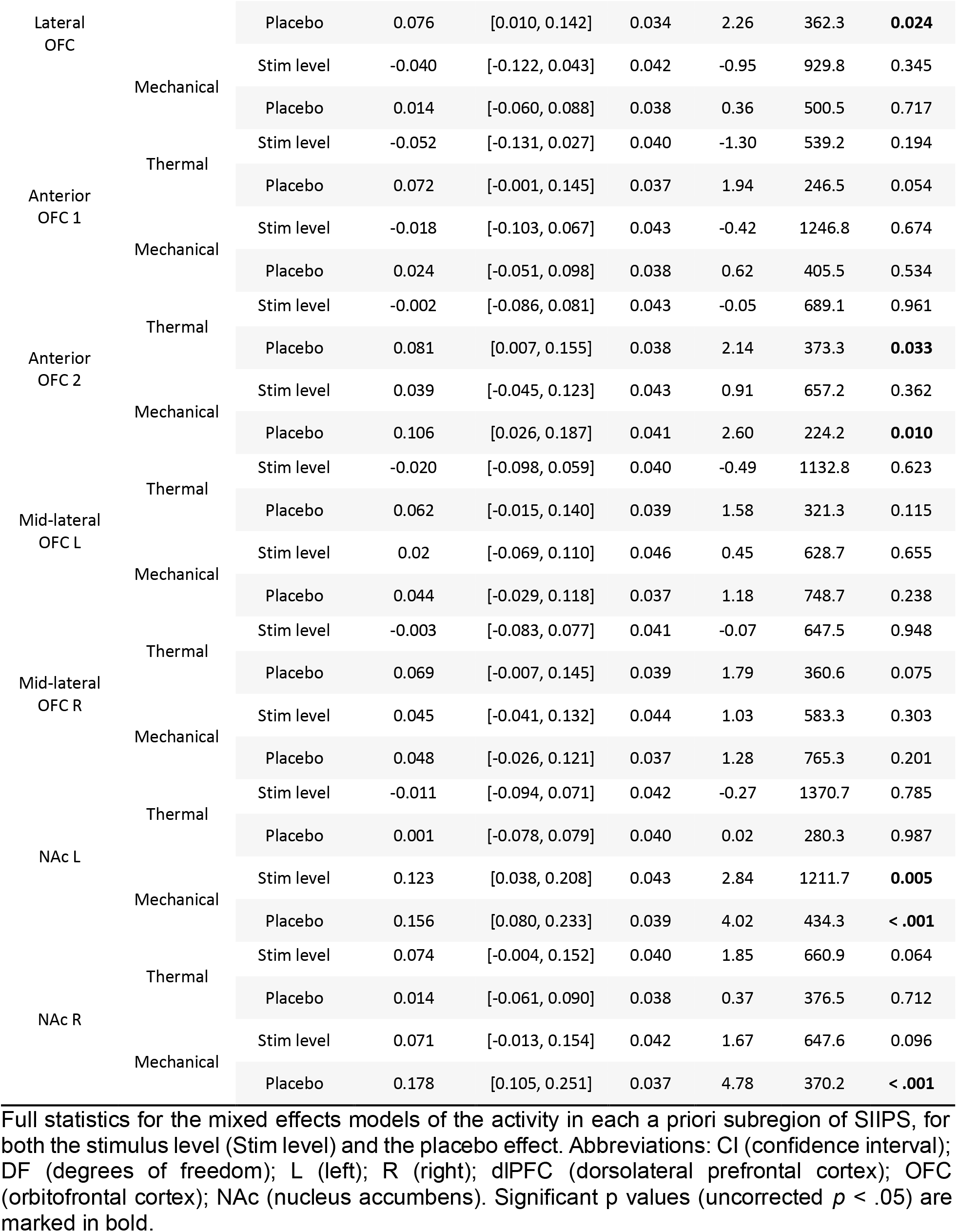
Statistics for a priori ROIs for higher-level pain processing: stimulus level and <u>placebo effects</u>.

### Individual differences in placebo effects

In the current paper, we have focused primarily on the causal effects of placebo treatment in experimental settings on pain at the group level, as reported above. Nevertheless, we also tested the correlations between individual differences in placebo analgesia and neural placebo-induced changes, as was done in previous studies. Importantly, correlations in small samples (as in most previous placebo fMRI studies) are unreliable, and meta-analyses cannot address this issue because of heterogeneity across studies. Thus, this is one of the first studies which could adequately test these behavioral-neural correlations. Nevertheless, such correlations do not imply causal effects of placebo, and can be driven by other processes (see Discussion).

#### NPS

Although there was no effect of placebo treatment on the NPS on average at the group level, we found that stronger placebo-induced NPS reductions were associated with stronger behavioral analgesia in both the thermal and mechanical modalities (mixed-effects model, see methods for details; thermal: β = 0.198, *SE* = 0.034, *t*_(106.2)_ = 5.83, *p* < .001, 95% CI = [0.131, 0.265]; mechanical: β = 0.202, *SE* = 0.037, *t*_(63)_ = 5.46, *p* < .001, 95% CI = [0.128. 0.276]; Figure 6A). Since different participants use the rating scale differently, increasing skewness in the data, we tested the robustness of these results by testing Spearman’s correlations based on the rank-order of the behavioral and neural Placebo – Control values across participants (ranked separately within each combination of pain modality and stimulus level). The correlation was significant with this nonparametric correlation test as well (thermal: β = 0.158, *SE* = 0.033, *t*_(159.8)_ = 4.84, *p* < .001, 95% CI = [0.094, 0.222]; mechanical: β = 0.140, *SE* = 0.032, *t*_(258.4)_ = 4.43, *p* < .001, 95% CI = [0.078. 0.202]). Since there was no group effect of placebo treatment on the NPS, these findings could result from NPS reductions in stronger placebo responders, or from other correlated factors such as random differences in sensitivity between the Placebo and Control skin sites. We cannot identify placebo responders from independent data in the current study, and therefore we cannot dissociate between these two alternatives (see Discussion). NPS placebo-induced reductions were not correlated between the thermal and mechanical modalities (β = 0.067, *SE* = 0.056, *t*_(99.3)_ = 1.18, *p* = .239, 95% CI = [-0.045, 0.179]).

#### Nociceptive regions

We tested the correlation between the placebo-induced neural reductions and the behavioral analgesia in each of the a priori nociceptive ROIs (see Supplementary Table 5 for full statistics). In the thermal modality, the correlation was significantly positive in the aMCC, right dpIns, PAG and medial thalamus (the result in the medial thalamus did not survive Bonferroni correction). In the mechanical modality, we found significant positive correlations in the aMCC, right and left dpIns, and PAG. These indicate greater placebo-induced reduction in brain response with greater analgesia, as expected. Although the remaining regions did not show statistically significant differences, trends were all in the positive direction in both modalities.

**Figure 6.**
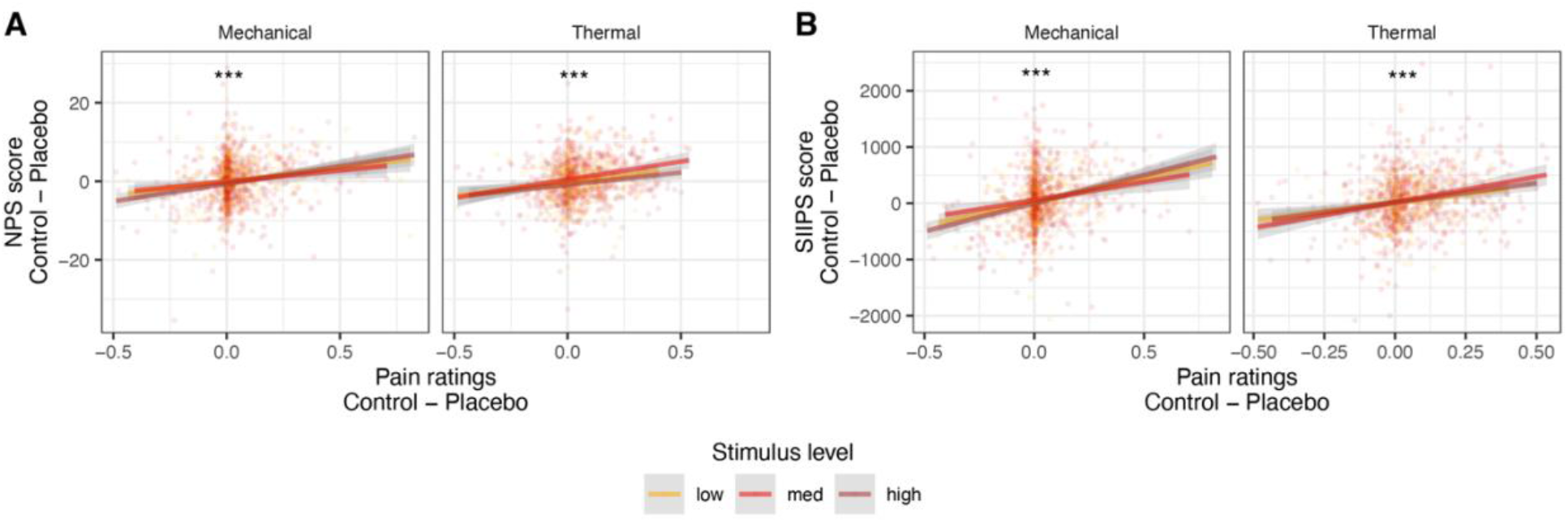
Correlations between behavioral and neural placebo-induced reductions. Placebo-induced neural reductions (Control – Placebo) in the (**A**) NPS or (**B**) SIIPS, as a function of the behavioral analgesia (pain ratings Control – Placebo) for each stimulus intensity level (color) and modality. Each dot is a participant. Lines represent linear smoother, with the 95% confidence interval shaded in gray. Asterisks represent significance of the correlation (uncorrected): * *p* < .05, ** *p* < .01, *** *p* < .001.

#### SIIPS

Placebo-induced neural reductions in SIIPS scores were significantly correlated with behavioral analgesia in both modalities, whether measured linearly (thermal: β = 0.229, *SE* = 0.033, *t*_(302.3)_ = 7.00, *p* < .001, 95% CI = [0.165, 0.294]; mechanical: β = 0.241, *SE* = 0.036, *t*_(67.9)_ = 6.78, *p* < .001, 95% CI = [0.170, 0.312]; Figure 6B) or based on ranks (Spearman; thermal: β = 0.207, *SE* = 0.032, *t*_(170.3)_ = 6.47, *p* < .001, 95% CI = [0.144, 0.270]; mechanical: β = 0.181, *SE* = 0.034, *t*_(177)_ = 5.27, *p* < .001, 95% CI = [0.113, 0.249]). Moreover, placebo-induced SIIPS reductions positively correlated with placebo-induced NPS reductions in both modalities (thermal: β = 0.243, *SE* = 0.033, *t*_(243.1)_ = 7.32, *p* < .001, 95% CI = [0.178, 0.309]; mechanical: β = 0.188, *SE* = 0.036, *t*_(173.5)_ = 5.21, *p* < .001, 95% CI = [0.117, 0.260]). Placebo-induced SIIPS reductions did not correlate across thermal and mechanical modalities (β = 0.081, *SE* = 0.065, *t*_(85.4)_ = 1.26, *p* = .212, 95% CI = [-0.047, 0.210]).

#### SIIPS subregions

The correlation between the behavioral and neural placebo-induced reductions in local pattern responses within SIIPS subregions was significant only for the right lingual gyrus (a positive correlation, as expected) in the thermal modality, and the left superior temporal gyrus and right temporal pole (both with negative correlations, surprisingly, however not surviving correction for multiple comparisons) in the mechanical modality (see Supplementary Table 6 for full statistics).

#### Higher level regions

In these a priori regions, we expected greater placebo-induced responses (more negative control – placebo scores) to predict stronger behavioral analgesia (control – placebo; a negative correlation). Nevertheless, we did not find significant negative correlations in any of these regions in both modalities. Conversely, in the thermal modality we found significant positive correlations in the right dlPFC and more anterior area of the left dlPFC, and in the right NAC (however only the effect in the right dlPFC survived Bonferroni correction), and in the mechanical modality we found a significant positive correlation only in the more anterior region of the dlPFC (not surviving Bonferroni correction; see Supplementary Table 7 for full statistics). Thus, placebo-induced activation predicted weaker placebo effects in dlPFC, anterior PFC, and NAC.

## Discussion

Previous studies have provided mixed evidence on nociceptive modulation in placebo analgesia, with small, individual studies showing evidence for opioid release ^42–46^ and spinal modulation ^40,41^, but meta-and mega-analyses revealing small effects of placebo in nociceptive regions. Here, in the largest neuroimaging study of placebo effects to date, we investigated both conditioned placebo effects in thermal pain (the most common placebo paradigm) and transfer to unconditioned placebo effects in mechanical pain, a novel test of generalization. Placebo effects on reported pain were highly significant, and effect sizes were moderate to large (d ∼= 0.6), comparable to those found in previous studies, and were as large in the mechanical pain transfer test as they were in the conditioned pain condition. In addition, placebo effects were modestly correlated across conditioned thermal and mechanical transfer conditions, but only correlated with pre-scan expectations in the thermal condition. These findings are important because how broadly placebo effects transfer across contexts and outcomes (e.g., pain responses) is a crucial and heavily debated issue ^80–84,86,87,89–91^. On one hand, nociceptive pathways are organized by stimulus type, with different channels in the periphery ^102,103^ and somewhat divergent cortical representations ^104,105^ and pain sensitivity ^106–109^, and placebo effects are largely uncorrelated across pain types in some previous studies ^80^ (though these are likely underpowered given the effect sizes we report here). On the other hand, recent behavioral evidence suggests that placebo effects based on associative learning often do transfer to different challenges to produce benefits ^83,89–91^ and in some cases harms ^86^. Our findings indicate robust behavioral placebo effects that are generalizable to new outcomes (i.e., a new pain modality). In addition, they show generalizability at the neural level for pain valuation and motivation-related brain processes, as we describe below.

Despite these substantial behavioral effects, fMRI analyses did not reveal significant reduction in the NPS–the most widely validated neuromarker of nociceptive pain to date–or nociception-related ROIs that were pre-registered based on previous studies. Bayes Factor analyses showed strong evidence in favor of null effects. Bayesian evidence for null effects was also found in individual subregions, precluding the possibility that the null effects resulted from a mix of positive and negative findings in different regions. In addition, in the novel test of transfer to mechanical pain, placebo treatment caused paradoxical activity increases in several of the regions most closely associated with pain processing, including aMCC, dpINS, and sensory thalamus (VPL/VPM). The NPS has been shown to be sensitive to bottom-up stimulus intensity across multiple pain types ^54^, including sensitivity to both thermal and mechanical pain intensity in this study, and our sample size provided high power to detect both small effects and provide strong Bayesian evidence in favor of the null. With n = 392 and p < 0.05, we have more than 80% power to detect “very small” effects of d = 0.15, and ∼100% power to detect “small” effects of d = 0.3. Thus, together, these findings suggest that placebo analgesia is not driven by modulation of low-level nociceptive processes, at least for the average participant in experimental settings.

However, pain is more than nociception, and incorporates substantial affective and evaluative contributions ^110^. We found evidence that placebo effects reduce activity in brain systems related to higher-level evaluative contributions to pain. The SIIPS neuromarker was trained to capture endogenous brain contributions to pain above and beyond stimulus intensity, reflecting these aspects. It is tempting to associate SIIPS with the affective aspects of pain, but strong correlations between pain affect and intensity measures were found in the current study and previous ones ^111–113^; thus, we believe SIIPS is most closely associated with evaluative and value-construction aspects of pain. Here, the SIIPS response to thermal pain was reduced with placebo treatment, and this effect transferred to unconditioned mechanical pain. Effects in local regions were generally smaller, suggesting that reductions were distributed across brain systems, but particularly prominent reductions were found in the mechanical pain condition in the NAc, STG, and TP. The NAc has shown activation in previous meta-analyses of placebo ^51,114^, placebo-induced opioid release ^45,46,115^ and dopamine release ^115^, placebo-induced changes in reward prediction error signaling ^116^, and mediation of cognitive reappraisal effects on pain ^117^. However, a recent meta-analysis showed evidence for NAc increases with pain but little evidence for placebo effects ^53^. It may be that placebo-induced NAc increases are masked in many studies because placebo simultaneously directly activates and reduces pain-related inputs to NAc; future studies must disentangle the effects of pain of different types as well as placebo (or other ‘top-down’ effects) to test this further. STG and TP have seldom been discussed in the placebo literature, but they are extensively involved in the construction of emotional experiences and emotional memories ^118^.

In addition to effects on SIIPS, we found placebo-induced activity in dlPFC, anterior PFC, and OFC. These areas were suggested as key mediators of placebo effects in early studies ^49^ but effects were not consistent across studies in a recent meta-analysis ^53^. Together, findings on the SIIPS, regions associated with high-level pain evaluation and construction of emotion, and prefrontal cortical increases support the conclusion that placebo analgesia is mostly driven by higher-level neural processing, including regions that are related to the construction of value and motivation.

The paradoxical placebo-induced increases in mechanical pain responses in some pain-related regions–aMCC, dpINS, and thalamus–warrant further discussion. Placebo-induced increases have not been reported in previous, smaller-scale studies to our knowledge, though these studies have not focused on transfer to unconditioned pain types. One possibility is that these increases reflect aversive prediction errors, an account broadly consistent with the predictive coding framework, with BOLD signals reflecting prediction errors from lower-level processing stages (i.e., pain-related signals that are not “canceled out” by predictions) ^21,24^. Roy et al., 2014 ^119^ and Geuter et al., 2017 ^71^ provided some evidence for aversive prediction error coding in INS and aMCC, and Roy et al. also found a paradoxical placebo-related increase in PAG. Combined with our findings of self-reported placebo analgesia, these findings suggest a dissociation between behavioral and neural correlates of pain. While pain reports integrate top-down predictions provided by the placebo context (i.e., assimilation effects), consistent with Bayesian accounts of placebo effects ^24^, nociceptive systems show the opposite effects (i.e., contrast effects), consistent with bottom-up signals that are not canceled out by predictions (because actual nociceptive input is higher than predicted under placebo treatment). This account is consistent with placebo effects on later-stage decision-making and experience construction rather than on early nociceptive signals.

This account is also consistent with a dual-process theory of pain modulation, in which safety cues that reduce the threat value of pain can have dual, opposing effects. On one hand, safety cues could allow competing motivational processes to gain priority and suppress pain, mediated in part by “motivation/decision” circuits like the NAc and medial PFC ^120–122^. On the other hand, threat of pain induces preparatory processes that also suppress nociception (threat– and stress-induced analgesia), including endogenous opioid release ^123–125^. Safety cues that reduce threat would block this effect. If threat analgesia has effects on lower levels of the nociceptive processing hierarchy than safety-related expectancies, placebo effects would increase nociceptive input (by reducing threat analgesia) while reducing pain valuation and motivation (in NAc/MPFC). Brain regions with a strong sensory component, like dpINS, sensory thalamus, and pain-selective portions of aMCC ^126,127^, would show paradoxical placebo-related increases such as those we observe here. Since these placebo-induced increases in nociceptive pain regions were only found in the transfer condition (i.e., in mechanical but not thermal pain trials), they may result from a transfer of the threat analgesia but no (or weaker) transfer of the safety-related expectancies.

Finally, this study provided a rare opportunity to examine individual differences in neural placebo effects in a sufficiently large sample to allow for stable correlation estimates and detection of small effects typical of between-person correlations ^128,129^. We found significant correlations between behavioral analgesia and larger reductions in both the NPS and SIIPS, as well as some individual regions. This suggests that placebo treatment may cause reductions in the NPS in some individuals, as suggested by a previous person-level meta-analysis ^54^. However, we cannot endorse this conclusion here because such correlations cannot provide strong evidence for causal effects. Here, as in previous literature, the strength of an individual’s behavioral analgesic effect is conflated with random variation in sensitivity on different skin sites and/or sensitization/habituation over time. This issue is common to virtually all clinical trials of treatment effects. Future studies could productively examine brain-behavioral correlations by selecting high and low placebo responders based on independent criteria (e.g., a separate session using different skin sites). Here, pre-scan expectation ratings could serve this goal, but expectancy ratings were not correlated with neural placebo-control differences, and were only associated with behavioral analgesia in thermal pain.

Individual difference correlations across thermal and mechanical pain do not suffer from this problem (because different skin sites were used for each modality). However, these correlations were significant behaviorally but non-significant for both NPS and SIIPS, suggesting low trait-like stability of neural placebo responses. This is important because the ability to predict who will be a placebo responder is a crucial and perennial issue in medicine–with many attempts to predict and control for placebo effects in drug and device trials ^130,131^– and such predictions hinge on the ability to identify stable traits across outcomes and time. Whereas some studies have found correlations between placebo effects and trait-like measures including genetics ^132–134^, brain structure and personality ^135,136^, these studies have been small and underpowered for the data types and expected effect sizes ^128,137^, and a recent meta-analysis of personality shows inconsistent effects across studies ^82^. One of our initial hypotheses was that while self-reported placebo effects and personality may be unstable across contexts (e.g., ^81^), neural responses would be more stable at the individual differences level and show stronger correlations across pain types, but this did not appear to be the case here.

There are several additional limitations to the current study that should be taken into account. First, the overall pain level across participants was relatively low based on the subjective pain ratings, with the vast majority of ratings indicating less than a moderate amount of pain, particularly in the mechanical modality. These may have been the result of the specific stimulus intensities chosen, or of using the finger as the body site of stimulation, since glabrous skin lacks type II a delta fibers ^138^. Nevertheless, we observed strong placebo effects of the ratings, as well as substantial variability across participants that allowed us to test for individual differences. Second, the use of an imaging protocol with a multiband acceleration factor of 8 may have enhanced signal dropout in some areas, such as ventral prefrontal areas and NAc ^139^, though we did find effects in some of these regions. Third, we used the canonical hemodynamic response function (cHRF), as was pre-registered and done in most previous studies. However, early studies ^43,49^ have distinguished between early and late responses without assuming cHRF, and different HRF models may yield different results. Fourth, different placebo-induction protocols or analysis pipelines may lead to different findings ^140^, for example with regard to the effect of placebo on nociceptive processes. Here, the placebo induction combined several components in order to maximize placebo effects (e.g., suggestion, conceptual conditioning and classical conditioning), which yielded strong behavioral effects, in addition to an unconditioned transfer condition (which produced similarly large behavioral and neural effects). In addition, the analysis pipeline used is a common one and many of his components were pre-registered.

Overall, addressing the critical question of the neural level at which placebo manipulations operate, our findings strongly indicate that a combination of conditioning and suggestion leads to large reductions in reported pain that are manifested in neural changes in higher-level brain regions related to pain, decision-making and motivation, but not lower-level nociceptive pain regions. These effects are not limited to conditioned effects, but transfer to outcomes that utilize at least partially distinct neural circuitry. More broadly, our findings point at the importance and promise of new treatments that are based on beliefs and expectations, in pain and beyond, that operate on higher-level neural processes that could yield meaningful clinical improvements ^141,142^. Such new treatments are critically needed now even more than ever, as pain remains an essential but poorly understood motivating force and chronic pain remains undertreated. Continued studies of placebo effects are needed to understand how the brain constructs pain experience and how those experiences drive our long term experience and behavior.

## Methods

### Participants

Participants were recruited by telephone from the Colorado Community Twin Sample, which is derived from the Colorado Twin Registry, a population-based registry which has been run by the Institute of Behavioral Genetics (IBG) at the University of Colorado since 1984 ^143^. The study includes a larger sample of participants, but this study is based on a pre-registered dataset of 397 participants who completed their participation and whose data were preprocessed by September 1st, 2022, when we pre-registered the study (pre-registration link: https://osf.io/unh7f). Participants were excluded from participation in the study if they did not pass MRI screening (e.g., add metals in their body) or had a history of liver disease/damage, allergies to local analgesics, or were breastfeeding.

Some participants have been excluded prior to the pre-registration due to a protocol change early in the study (n = 11), data loss/corruption (n = 11), errors related to our thermal stimulator (n = 1) or inconsistencies in the placebo induction/treatment protocol (n = 3). Two participants who were included in the pre-registered sample were excluded from analyses because of corrupted behavioral data. The final dataset included 395 participants, of which 142 are monozygotic twins, 160 are dizygotic twins, and 93 are individuals without siblings in the dataset. Participants’ age ranged from 30 to 43 years (*M* = 35.43, *SD* = 2.60), and the sample included 164 men and 231 women. All participants provided their informed consent at the beginning of the experiment. The experiment was approved by the institutional review board of the University of Colorado Boulder.

Individual stimulus trials were excluded if pain intensity rating response times were above 5.01 or below 0.02 seconds. These indicate either that the participant did not respond within the time allotted (five seconds) or responded too quickly to represent deliberate ratings. Entire participants were dropped if after dropping trials or scans due to above exclusion criteria, they lacked complete sets of conditions (placebo-thermal, placebo-mechanical, control-thermal, control-mechanical) for more than one stimulus level.

### Procedures

First, participants were presented with an overview of the experiment components and completed a short survey about their health and mood. Then, they completed an anti-saccade task, provided saliva samples, practiced the rating scale, and completed calibration tasks. The rating scale was a generalized Labeled Magnitude Scale (gLMS; ^96^), with anchors of “no pain” or “no unpleasantness” and “most intense sensation imaginable” or “most unpleasantness imaginable”. The calibration tasks were used to ensure participants’ tolerance to the painful thermal stimuli. Stimuli in the calibration task ranged between 45.5 – 48.5 °C, with a duration of 10 seconds each. Participants rated each stimulus verbally. Participants who were unable to tolerate the stimuli were excluded from further participation. The thermal stimuli were delivered with a 16 × 16 mm surface thermode (PATHWAY ATS; Medoc, Inc, Israel).

Following the calibration task, the placebo manipulation took place. Participants were given two identical creams with different instructions. One cream was introduced as “Prodicaine, an effective pain-relieving drug” (the placebo cream), and the second cream was introduced as “a control cream with no effects” (the control cream). These two creams were applied to two different fingers of the left hand of the participant (cream allocation to fingers was counterbalanced across participants, and the cream was applied to the entire inner part of each finger). Participants watched a video describing the prodicaine application procedure and a short testimonial from an (allegedly) pilot participant. They also received forms disclosing potential side effects of the Prodicaine, with realistic-looking drug company logos, in a room with medical equipment and related contextual cues.

To strengthen expectations of pain relief from the placebo treatment cream, participants were subjected to two conditioning protocols. Importantly, both tasks were based on thermal, not mechanical, stimuli. First a “symbolic” conditioning paradigm was administered: an inert thermode was placed on the control site and the participants completed a trial sequence mimicking a thermal stimulation sequence, except instead of thermal stimuli they were shown ratings they were told come from prior participants. These ratings were systematically high for 16 stimuli. The thermode was then moved to the prodicaine treated site and the procedure was repeated for 32 stimuli. This time the ratings were systematically low. Finally the thermode was moved back to the control site and 16 additional ratings were once again systematically high. The thermode was placed on the proximal phalanges. Second, a classical condition paradigm was administered. This paradigm was identical to the symbolic conditioning paradigm, except instead of being shown ratings of other participants, participants were subjected to noxious thermal stimulation of the proximal phalanges. They were told that stimuli were all of the same intensity, but the intensity was surreptitiously lowered by 3.5 °C when stimulating the prodicaine treated site compared to the control treated site (44 and 44.5 °C for the placebo cream and 47.5 and 48 °C for the control cream). Following each stimulus, participants rated the intensity and unpleasantness of the stimulus. Following the conditioning task, participants rated their expectations regarding the Prodicaine efficacy in the next task, on a linear scale between 0 [not at all] to 100 [most effective].

Following the placebo manipulation, the test phase was conducted in the MRI scanner. The creams were reapplied to four fingers (index, middle, ring, pinky), two of which to be used for thermal and two for mechanical (unconditioned) stimuli, with one finger for the placebo cream and one for the control cream in each modality (fingers allocation was counterbalanced across participants). Mechanical pain stimuli were administered using an in-house pressure pain device, an MRI-safe device with dynamic pressure delivery controlled by LabView (National Instruments). The test task included 32 stimuli divided into four runs, with four thermal and four mechanical stimuli in a random order per run. Three stimulus intensities were used for each modality: low, medium, and high (thermal: 46.5, 47 and 47.5 °C; mechanical: levels 3,4,5, corresponding to about 6, 7 and 8 kg/cm^2^, respectively). Half of the stimuli from each modality were delivered to the finger with the control cream, and half to the finger with the placebo cream, such that the first and last run included stimulation of the control cream skin site, and the second and third included stimulation of the placebo cream skin site (a control-placebo-placebo-control design). Each stimulus lasted 10 seconds, and was delivered to the distal phalanges of the left hand. Before each stimulation, an anticipatory cue was presented for 0.5 second followed by a one or three seconds randomly-jittered delay. Three seconds after each stimulus, participants rated its intensity and unpleasantness (five seconds each, in a random order) with a tracking ball.

#### Imaging data acquisition

Imaging data were acquired using a 3T Siemens Prisma MRI scanner with a 32-channels head coil, at the University of Colorado at Boulder. First, a T1-weighted structural scan was acquired using a magnetization prepared rapid gradient echo (MPRAGE) pulse sequence with parallel imaging factor (iPAT) of 3, TR = 2000 ms, TE = 2.11 ms, flip angle = 8 degrees, FOV = 256 mm, resolution = 0.8 × 0.8 × 0.8 mm. Then, we acquired fieldmaps for each participant, one with a posterior-anterior (PA) and one with an anterior-posterior (AP) direction, with the following imaging parameters: TR = 7220 ms, TE = 73 ms, flip angle = 90 degrees, FOV = 220 mm, and in plane resolution of 2.7 × 2.7 × 2.7 mm. Resting-state fMRI data was acquired using T2*-weighted echo-planar imaging (EPI) sequence with multiband acceleration factor of 8, TR = 460 ms, TE = 27.20 ms, flip angle = 44 degrees, FOV = 220 mm, resolution of 2.7 × 2.7 × 2.7 mm, 56 slices, and 816 acquired volumes. Then, participants completed four runs of the pain test task while scanned with a similar fMRI sequence, obtaining 550 volumes for each of the four task fMRI runs. They then completed a perspective-taking fMRI task (similar protocol, 1322 volumes; this task is outside the scope of the present paper). Finally, two diffusion-weighted (dMRI) scans were acquired, one with a PA and one with an AP acquisition direction. The dMRI PA scan included a multiband acceleration factor of 3, 47 directions, TR = 4000 ms, TE = 77 ms, FOV = 224 mm, flip angle = 84 degrees, with a b-value of 2400 s/mm^2^. The AP dMRI scan was similar, except for an inverse acquisition direction and 44 diffusion directions acquired.

#### Imaging data preprocessing

Structural and functional data were preprocessed using fMRIPrep version 20.2.3 ^144^ (RRID:SCR_016216), which is based on *Nipype* 1.6.1 (RRID:SCR_002502; ^145,146^).

##### Anatomical data preprocessing

The T1-weighted (T1w) image was corrected for intensity non-uniformity (INU) with N4BiasFieldCorrection ^147^, distributed with ANTs 2.3.3 (RRID:SCR_004757; ^148^), and used as T1w-reference throughout the workflow. The T1w-reference was then skull-stripped with a *Nipype* implementation of the antsBrainExtraction.sh workflow (from ANTs), using OASIS30ANTs as target template. Brain tissue segmentation of cerebrospinal fluid (CSF), white-matter (WM) and gray-matter (GM) was performed on the brain-extracted T1w using fast (FSL 5.0.9, RRID:SCR_002823; ^149^). Brain surfaces were reconstructed using recon-all (FreeSurfer 6.0.1, RRID:SCR_001847; ^150^), and the brain mask estimated previously was refined with a custom variation of the method to reconcile ANTs-derived and FreeSurfer-derived segmentations of the cortical gray-matter of Mindboggle (RRID:SCR_002438; ^151^). Volume-based spatial normalization to two standard spaces (MNI152NLin2009cAsym, MNI152NLin6Asym) was performed through nonlinear registration with antsRegistration (ANTs 2.3.3), using brain-extracted versions of both T1w reference and the T1w template. The following templates were selected for spatial normalization: *ICBM 152 Nonlinear Asymmetrical template version 2009c* (RRID:SCR_008796; TemplateFlow ID: MNI152NLin2009cAsym; ^152^], *FSL’s MNI ICBM 152 non-linear 6th Generation Asymmetric Average Brain Stereotaxic Registration Model* (RRID:SCR_002823; TemplateFlow ID: MNI152NLin6Asym ^153^).

##### Functional data preprocessing

The single-band reference (SBRef) was used as a reference volume along with its skull-stripped version. A B0-nonuniformity map (or *fieldmap*) was estimated based on two EPI references with opposing phase-encoding directions, with 3dQwarp (AFNI 20160207; ^154^). Based on the estimated susceptibility distortion, a corrected EPI reference was calculated for a more accurate co-registration with the anatomical reference. The BOLD reference was then co-registered to the T1w reference using bbregister (FreeSurfer) which implements boundary-based registration ^155^. Co-registration was configured with six degrees of freedom. Head-motion parameters with respect to the BOLD reference (transformation matrices, and six corresponding rotation and translation parameters) are estimated before any spatiotemporal filtering using mcflirt (FSL 5.0.9, ^156^). First, a reference volume and its skull-stripped version were generated using a custom methodology of *fMRIPrep*. The BOLD time-series were resampled onto the following surfaces (FreeSurfer reconstruction nomenclature): *fsnative*, *fsaverage6*, *fsaverage*. The BOLD time-series were resampled onto their original, native space by applying a single, composite transform to correct for head-motion and susceptibility distortions. These resampled BOLD time-series will be referred to as *preprocessed BOLD in original space*, or just *preprocessed BOLD*. The BOLD time-series were resampled into standard space, generating a *preprocessed BOLD run in MNI152NLin2009cAsym space*. First, a reference volume and its skull-stripped version were generated using a custom methodology of *fMRIPrep*. All resamplings can be performed with *a single interpolation step* by composing all the pertinent transformations (i.e. head-motion transform matrices, susceptibility distortion correction when available, and co-registrations to anatomical and output spaces). Gridded (volumetric) resamplings were performed using antsApplyTransforms (ANTs), configured with Lanczos interpolation to minimize the smoothing effects of other kernels ^157^. Non-gridded (surface) resamplings were performed using mri_vol2surf (FreeSurfer). Following the preprocessing with fMRIPrep, data were smoothed with a Gaussian kernel of 6 mm.

## Data analysis

### Participant-level analysis

The first level model included the following regressors: four regressors for the cue period (one for each combination of modality [thermal / mechanical] and condition [placebo / control]), 12 regressors for the pain-evoked period, and a regressor for the rating period. We modeled 12 experimental conditions for the pain-evoked period in a 3 x 2 x 2 factorial design–including Stimulus level (3 levels), Modality (2 levels, thermal and mechanical), and Placebo treatment (2 levels, placebo and control)–using a separate regressor for each condition. Indicator vectors ([0,1]) indicating the presence or absence of each condition were convolved with a canonical hemodynamic response function implemented by spm12. Nuisance regressors included 24 motion regressors (six motion parameters–translation and rotation in three directions–together with their derivatives, quadratics and derivatives of the quadratics) and a mean CSF signal regressor (estimated by fMRIPrep during preprocessing). We also implemented spike censoring, with spikes identified by our in-house spike detection algorithm implemented by CanlabCore/diagnostics/scn_session_spike_id.m (available at github.com/canlab/CanlabCore). The first eight frames of each run were censored, and a 180 Hz high pass temporal filter was implemented using cosine basis functions. This procedure resulted in 12 separate parameter estimate maps for each stimulus condition (e.g., high intensity thermal stimuli delivered to the placebo skin site), for each participant.

Statistical analysis was performed using Matlab 2020a and CANlab neuroimaging analysis tools (shared via Github at https://canlab.github.io/; the version used was from the beginning of August 2022, with the last commit of Canlab Core being b6db85e, SHA b6db85e2577d967d90c3bbe508c3c6acff37e268). Activity in each a priori region of interest or neuromarker for each contrast (each condition of each participant) image was computed as a continuous score. For a priori regions of interest, the score was based on the averaged univariate GLM-derived BOLD contrast estimates across voxels within the region of interest. For neuromarkers and their subregions, the score was based on the dot product of the univariate map with the neuromarker weight map.

The different scores, as well as the behavioral pain ratings, were tested at the group level using a mixed effects model. In the case of datasets with twins, treating individuals as independent observations leads to inflated estimates of degrees of freedom and consequently overly liberal statistical inference. To account for this, we introduce familial random effects. Thus, fixed effects can be interpreted as the mean effect taken over families rather than individuals. Twin studies additionally introduce heteroscedastic sources of variance since, within families, dizygotic twins are expected to be more variable than monozygotic twins. Naive maximum likelihood methods would produce fixed effect parameter estimates that are biased towards the dizygotic twins in the sample. To prevent that, we instead model each category of twin using separate random effects covariance parameters. Such a model can be implemented using off the shelf mixed effects modeling software with an appropriate covariance model specification and coding scheme.

We used Matlab’s fitlme function to estimate effects of the placebo condition and the stimulus intensity. Formally, using Wilkinson notation ^158^:

score ∼ 1 + stimulus_level + placebo_condition + (1 + stimulus_level + placebo_condition | family_ID) + (intercept_monozygotic + stimulus_level_monozygotic + placebo_condition_monozygotic –1 | participant_ID)

+ (intercept_dizygotic + stimulus_level_dizygotic + placebo_condition_dizygotic

– 1 | participant_ID)

Where the stimulus_intensity was coded as –0.5, 0, 0.5 for low, medium, high, respectively, and the placebo_condition coded as –0.5 for control and 0.5 for placebo. All regressors with the suffix “_dizygotic” were 0 for monozygotic twins, and all regressors with the suffix “_monozygotic” were 0 for dizygotic twins. The random effects covariance matrix was also constrained to be block diagonal by zygosity, such that the covariances of parameters with the suffix “_monozygotic” and with the suffix “_dizygotic” are zero. This generalizes a heteroskedastic twin error model ^159^ to within participant repeated measures designs.

Each model was tested with the interaction term (between the stimulus level and placebo condition) as a fixed effect, and then without the interaction term if the interaction was not significant. In such a case, to increase interpretability, the mean effects reported in the main text were based on the model without the interaction. Note that the interaction was not significant for all tested neuromarkers and regions of interest. Data from each pain modality were tested separately. Subjective pain ratings and brain scores within each modality were z-scored. Satterthwaite’s method ^160^ was used to estimate degrees of freedom for the mixed effects models. As pre-registered, participants without full data for at least two stimulus levels were excluded from the analysis. This led to the exclusion of 22 trials (0.47% of total number of trials), from three participants. Overall, 392 participants were included in the models for the effects of placebo and stimulus intensity. When the effect size is presented in the text as Cohen’s d, it is based on the ratio between the mean / sd across participants, without accounting for the familial structure.

To test the correlation between the behavioral and neural placebo-induced reductions, we computed the corresponding analgesia value as the averaged pain rating or brain score for each participant in each stimulus modality and intensity level separately for the placebo and the control condition. We then computed the difference between these scores (control minus placebo) within each combination of modality and stimulus level for each participant, and z-scored the values across participants within each modality. We also computed the rank order of differences (not z-scored) within each combination of stimulus level and modality.

These values were inputted to the following mixed-effects model, computed again with fitlme in Matlab 2020a, separately for the thermal and mechanical pain modalities:

brain_score ∼ 1 + stimulus_level + behavioral_analgesia + (1 + stimulus_level

+ behavioral_analgesia | family) + (stimulus_level_monozygotic + behavioral_analgesia_monozygotic + intercept_monozygotic –1 | participant_ID)

+ (stimulus_level_dizygotic + behavioral_analgesia_dizygotic + intercept_dizygotic –1 | participant_ID)

Note that we report results based on p < .05, without correcting for multiple comparisons. This is because in the current paper there is a need to balance between type 1 and type 2 errors, as we are testing multiple a priori (and mostly pre-registered) brain signatures and regions of interest based on previous literature with a new, substantially larger sample. To be slightly more conservative, we further note when a specific result does not survive Bonferroni correction for multiple comparisons within a specific set of regions (i.e., within the nociceptive regions, within SIIPS subregions, or within the set of higher-level regions).

Bayes Factor analysis was performed using R version 4.2.2., with the BayesFactor package version 0.9.12-2. Bayes factors were computed as the ratio of evidence in favor of the alternative hypothesis, which is the full model including main fixed and random effects (without random slopes), and the null hypothesis, which is the same model without the fixed effect of interest (e.g., the placebo effect ^98^). For the prior distribution, we used a wide Cauchy prior distribution (rscale value of 1). We further tested the robustness of the Bayes Factor results to different scaling factors of the prior distribution (√2/2, 1, and √2) and to the inclusion of an interaction term in the alternative model.

In addition, we tested the correlations between pre-scan expectations of Prodicaine efficacy and placebo effects on the behavioral pain ratings, NPS score, and SIIPS score. The correlations were tested with a mixed-effects model similar to the one described above for the correlations between behavioral analgesia and placebo-induced reductions in brain responses, with the participant-level expectations as a predictor instead of the behavioral analgesia. We again tested these correlations separately for the thermal and mechanical modalities, and the expectations were z scored within each modality. Correlations between placebo effects in the thermal and mechanical modalities were computed with a mixed-effects model predicting placebo-induced analgesia (or placebo-induced reductions in NPS or SIIPS score) in the mechanical condition based on the placebo-induced analgesia in the thermal condition (with the full random effects structure at the family and participant level as in the other models described above).

### Pre-registration

Pre-registration was performed following preprocessing of the imaging data, but prior to performing the data analysis that informs the hypothesized outcomes. The behavioral analyses were not pre-registered, as the pre-registration focused on the neural predictors and correlates of placebo analgesia, assuming a behavioral effect of placebo analgesia as has been repeatedly shown in numerous previous studies. In addition, the pre-registration did not include the NPS tests. This is because the NPS analysis was already run on some of the participants prior to the submission of the pre-registration, and therefore could not have been considered a non-tested hypothesis like the other hypotheses. However, the test of the placebo effect on the NPS score was run on the full pre-registered sample only after the pre-registration was submitted. The Bayes Factor analysis was also not pre-registered, but was used to quantify evidence in favor of the null vs. alternative hypothesis, particularly when null hypothesis significance testing (NHST) results were not significant and testing potential evidence in favor of the null was important. Finally, the pre-registration included a long list of additional hypotheses regarding a priori neural signatures and brain patterns, as well as a comparison of early vs. late activity during pain period, and tests of heritability of effects. These analyses are beyond the scope of the current paper, and will be executed and published in future papers.

## Data and code availability

The behavioral and imaging data are shared on OpenNeuro: ds004746; doi:10.18112/openneuro.ds004746.v1.0.0. Analysis codes are publicly shared on Github: https://github.com/rotemb9/paingen-placebo-fmri-paper (release 1.0.0). CANlab neuroimaging analysis tools that were used as part of the analysis are available at https://canlab.github.io/ (see methods for the version used).

## Acknowledgements

Rotem Botvinik-Nezer is an Awardee of the Weizmann Institute of Science – Israel National Postdoctoral Award Program for Advancing Women in Science. The study was funded by a National Institutes of Health (NIH) grant (DA046064, PIs Tor Wager and Naomi Friedman). The funders had no role in study design, data collection and analysis, decision to publish or preparation of the manuscript. The authors thank Mattew Keller for helpful discussions. Thanks to Gordon Matthewson, Dan Kusko, Dan Ryan, Patricia Townsend, Clayton Schneider, Mickela Heilicher, Suebin Song, Abi Adams, Jia Moore and Alexa Gonzalez for helping with data collection.

## Supplementary Information

### Supplementary Results

#### Behavioral results with unpleasantness ratings

Results were quantitatively similar when using the unpleasantness rather than the intensity ratings provided by the participants after each trial. Participants’ unpleasantness ratings following noxious stimuli were significantly lower in the Placebo compared to the Control condition (thermal: Placebo *M* = 0.109, Control *M* = 0.162, β = –0.375, *SE* = 0.039, *t*_(234.8)_ = –9.62, *p* < .001, 95% CI = [-0.452, –0.298]; mechanical: Placebo *M* = 0.095, Control *M* = 0.130, β = –0.233, *SE* = 0.039, *t*_(363.5)_ = – 5.77, *p* < .001, 95% CI = [-0.298, –0.147]). As expected, unpleasantness ratings increased with increasing stimulus intensity (thermal: low intensity *M* = 0.111, medium intensity *M* = 0.128, high intensity *M* = 0.168, β = 0.399, *SE* = 0.031, *t*_(209.4)_ = 12.70, *p* < .001, 95% CI = [0.337, 0.461]; mechanical: low intensity *M* = 0.100, medium intensity *M* = 0.109, high intensity *M* = 0.128, β = 0.214, *SE* = 0.033, *t*_(151.4)_ = 6.57, *p* < .001, 95% CI = [0.149, 0.278]). The Intensity X Placebo interaction was not significant (thermal: β = –0.100, *SE* = 0.053, *t*_(1253.1)_ = –1.87, *p* = .062, 95% CI = [-0.204, 0.005]; mechanical: β = 0.013, *SE* = 0.038, *t*_(1178.3)_ = 0.35, *p* = .724, 95% CI = [-0.061, 0.087]).

### Robustness to covariates and demographic effects

#### Pain ratings

The results indicating effects of the placebo condition and the stimulus intensity on pain ratings were robust to the inclusion of demographic covariates, sex and age. To test this, we ran the same models as described in the main text, with sex and age as additional covariates. Since age is the same within twin pairs, and sex is the same for the vast majority (except some opposite-sex dizygotic twins), age was only added as a fixed effect, and sex was added as a fixed effect and a random effect at the family level.

Pain ratings were significantly lower in the Placebo compared to the Control condition (thermal: β = –0.359, *SE* = 0.037, *t*_(235.5)_ = –9.72, *p* < .001, 95% CI = [-0.432, –0.286]; mechanical: β = –0.240, *SE* = 0.040, *t*_(273.3)_ = –6.06, *p* < .001, 95% CI = [-0.317, –0.162]) and significantly higher for higher stimulus intensity (thermal: β = 0.409, *SE* = 0.031, *t*_(347.8)_ = 13.22, *p* < .001, 95% CI = [0.348, 0.470]; mechanical: β = 0.206, *SE* = 0.031, *t*_(129.4)_ = 6.53, *p* < .001, 95% CI = [0.143, 0.268]). Males reported significantly higher pain compared to women in the mechanical (β = 0.187, *SE* = 0.086, *t*_(177.1)_ = 2.18, *p* = .031, 95% CI = [0.017, 0.356]), but not in the thermal (β = –0.033, *SE* = 0.075, *t*_(266.16)_ = –0.44, *p* = .661, 95% CI = [-0.180, 0.115]) modality, while age did not affect pain ratings in either modality (thermal: β = –0.009, *SE* = 0.037, *t*_(326.9)_ = –0.23, *p* = .816, 95% CI = [-0.082, 0.065]; mechanical: β = 0.058, *SE* = 0.040, *t*_(207.0)_ = 1.42, *p* = .156, 95% CI = [-0.022, 0.137]).

Furthermore, we did not find significant sex differences in placebo analgesia for the thermal pain trials (Control minus Placebo, female: *M* = 0.054, *SD* = 0.103, *N* = 217; male: *M* = 0.053, *SD* = 0.102, *N* = 150; mixed effects model predicting placebo analgesia with fixed and random effects of stimulus level and sex, and a fixed effect of age: β = 0.030, *SE* = 0.081, *t*_(263.49)_ = 0.42, *p* = .706738, 95% CI = [-0.125, 0.194]). The placebo effect was also not significantly related to age (β = 0.073, *SE* = 0.040, *t*_(263.87)_ = 1.81, *p* = .072, 95% CI = [-0.006, 0.153]. These two effects were similarly not significant in the mechanical pain modality (sex: female: *M* = 0.0.39, *SD* = 0.121, *N* = 216; male: *M* = 0.041, *SD* = 0.132, *N* = 152; β = –0.014, *SE* = 0.094, *t*_(237.41)_ = –0.15, *p* = .883, 95% CI = [-0.198, 0.171]; age: β = –0.021, *SE* = 0.046, *t*_(302.87)_ = –0.46, *p* = .649, 95% CI = [-0.111, 0.069]).

#### NPS

Including gender and age as covariates did not change the NPS results. The NPS score was significantly higher for higher intensity levels (thermal: β = 0.244, *SE* = 0.041, *t*_(678.9)_ = 5.99, *p* < .001, 95% CI = [0.164, 0.324]; mechanical: β = 0.163, *SE* = 0.041, *t*_(698.6)_ = 3.98, *p* < .001, 95% CI = [0.083, 0.243]) and was not significantly different as a function of the placebo condition (thermal: β = –0.031, *SE* = 0.036, *t*_(225.5)_ = –0.86, *p* = .329, 95% CI = [-0.102, 0.040]; mechanical: β = 0.008, *SE* = 0.037, *t*_(277.74)_ = 0.21, *p* = .831, 95% CI = [-0.065, 0.061]). There was a significant main effect of sex, such that the NPS score was significantly higher for females in the thermal (β = –0.200, *SE* = 0.067, *t*_(336.6)_ = –2.98, *p* = .003, 95% CI = [-0.332, –0.068]) but not mechanical (β = –0.063, *SE* = 0.074, *t*_(187.7)_ = –0.85, *p* = .265, 95% CI = [-0.209, 0.083]) pain trials, and there was no effect of age on the NPS score in both modalities (thermal: β = 0.001, *SE* = 0.034, *t*_(289.1)_ = 0.04, *p* = .971, 95% CI = [-0.065, 0.067]; mechanical: β = 0.033, *SE* = 0.035, *t*_(269.7)_ = 0.93, *p* = .356, 95% CI = [-0.037, 0.102]). Furthermore, the placebo-induced NPS reductions still significantly correlated with the behavioral analgesia in both modalities when controlling for age and sex (thermal: β = 0.199, *SE* = 0.034, *t*_(101.9)_ = 5.90, *p* < .001, 95% CI = [0.132, 0.266]; mechanical: β = 0.199, *SE* = 0.037, *t*_(60.1)_ = 5.35, *p* < .001, 95% CI = [0.125, 0.274]).

#### SIIPS

Similarly to the behavioral and NPS results, the SIIPS results were robust to the inclusion of gender and age as covariates. When including these demographic covariates, the SIIPS score was significantly lower in the Placebo compared to the Control condition in both the thermal (β = –0.128, *SE* = 0.033, *t*_(440.5)_ = –3.86, *p* < .001, 95% CI = [-0.194, –0.063]) and mechanical (β = – 0.111, *SE* = 0.037, *t*_(272.8)_ = –2.98, *p* = .003, 95% CI = [-0.185, –0.038]) modalities. The SIIPS score also increased with stimulus intensity in both modalities (thermal: β = 0.180, *SE* = 0.038, *t*_(1050)_ = 4.73, *p* < .001, 95% CI = [0.105, 0.255]; mechanical: β = 0.188, *SE* = 0.040, *t*_(232.4)_ = 4.76, *p* < .001, 95% CI = [0.110, 0.266]). The placebo-induced SIIPS reductions significantly correlated with the behavioral analgesia in both modalities also when controlling for age and sex (thermal: β = 0.237, *SE* = 0.033, *t*_(248.3)_ = 7.08, *p* < .001, 95% CI = [0.171, 0.303]; mechanical: β = 0.237, *SE* = 0.035, *t*_(86.4)_ = 6.73, *p* < .001, 95% CI = [0.167, 0.307]). In addition, the SIIPS score was significantly higher for females compared to males in the thermal (β = –0.190, *SE* = 0.074, *t*_(251.6)_ = –2.58, *p* = .010, 95% CI = [-0.336, –0.045]) but not in the mechanical (β = –0.027, *SE* = 0.073, *t*_(192.1)_ = –0.37, *p* = .714, 95% CI = [-0.172, 0.118]) pain modality, and there was no effect of age on the SIIPS score in both modalities (thermal: β = –0.012, *SE* = 0.036, *t*_(272.2)_ = –0.33, *p* = .745, 95% CI = [-0.083, 0.059]; mechanical: β = –0.003, *SE* = 0.036, *t*_(231.1)_ = –0.09, *p* = .931, 95% CI = [-0.075, 0.068]). The placebo-induced SIIPS reduction was stronger for younger participants in the thermal (β = –0.070, *SE* = 0.032, *t*_(418)_ = –2.20, *p* = .029, 95% CI = [-0.132, –0.007]) but not in the mechanical (β = 0.018, *SE* = 0.033, *t*_(401.1)_ = 0.56, *p* = .576, 95% CI = [-0.046, 0.083]) pain modality, and there were no sex effects on the placebo-induced SIIPS reductions (thermal: β = – 0.062, *SE* = 0.065, *t*_(222)_ = –0.95, *p* = .341, 95% CI = [-0.190, 0.066]; mechanical: β = –0.104, *SE* = 0.068, *t*_(204.8)_ = –1.51, *p* = .132, 95% CI = [-0.239, 0.031]).

### Additional, non-pre-registered SIIPS subregions

In addition to the eight pre-registered subregions of SIIPS that are described in the main text, we also tested for effects in seven subregions that were not pre-registered but are of interest based on recent work. Six of these subregions–right nucleus accumbens (NAc), left and right dlPFC, right secondary somatosensory cortex (S2), right sensorimotor cortex (SMC) and left precuneus– are “suppressors”, meaning that they are more active for higher stimulus intensities, but activity is associated with lower pain reports (i.e., in the original data based on which the SIIPS signature was developed ^75^ they had significant positive estimates for the effect of stimulus intensity, but their weights in the SIIPS signature are negative). The last region–the vmPFC– is a mediator of pain, but with a negative sign: It de-activates with increasing stimulus intensity, and greater deactivation is associated with greater pain controlling for stimulus intensity.

For full statistics in these seven subregions for both modalities, see Supplementary Table 4. Note that activity in SIIPS subregions represents the local pattern response, and thus (as explained in the main text) more positive responses indicate more pain-related activity in all regions. Following the findings from the previous study ^75^, we expected a significant positive effect of the stimulus intensity in the six suppressor regions, and a significant negative effect of stimulus intensity in the vmPFC. We further expected a significant placebo effect in all these seven regions, such that the pattern response score would be lower in the Placebo compared to the Control condition.

In the vmPFC, we found the expected significant negative effects of the stimulus intensity (not surviving Bonferroni correction) and the placebo treatment only in the mechanical modality. These effects were not significant in the thermal modality. In the suppressor regions, the placebo effect was significant only in the right NAc and left precuneus, and only in the mechanical modality. As for the stimulus level effect, it was not significantly positive in any of the suppressor regions. It was, however, unexpectedly significantly negative in the right S2 for both modalities, and in the right SMC and right dlPFC (the latter not surviving Bonferroni correction) only for the thermal modality.

## Supplementary Figures

**Supplementary Figure 1.**
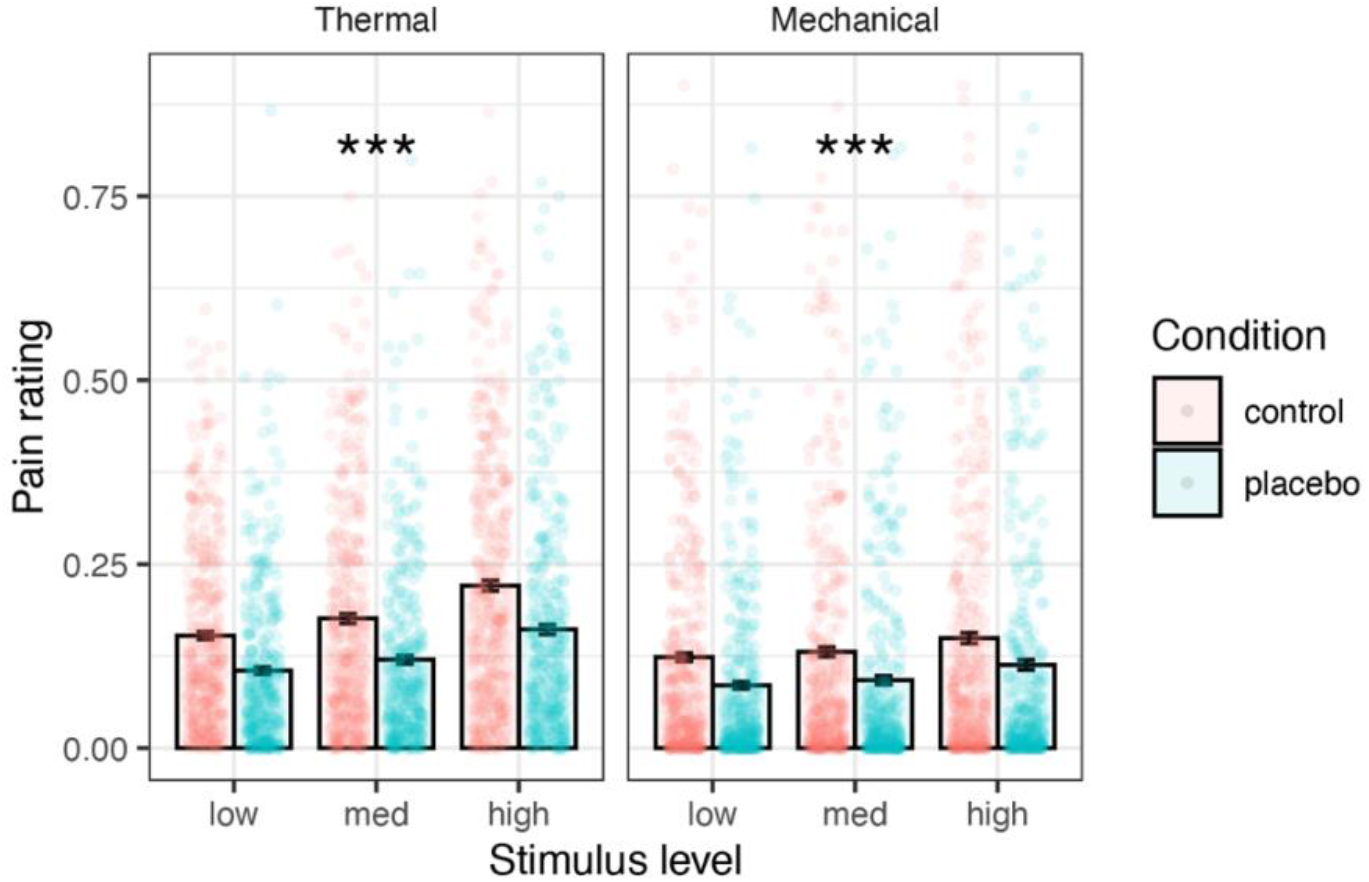
Behavioral results. The mean pain ratings across participants are presented, in the control (red) and placebo (blue) condition, for each combination of modality and stimulus level. Error bars represent within-participant standard error of the mean, based on Morey, 2008 ^97^. Points represent single participants. Asterisks represent significance of the placebo effect (Placebo vs. Control, uncorrected): * *p* < .05, ** *p* < .01, *** *p* < .001.

**Supplementary Figure 2.**
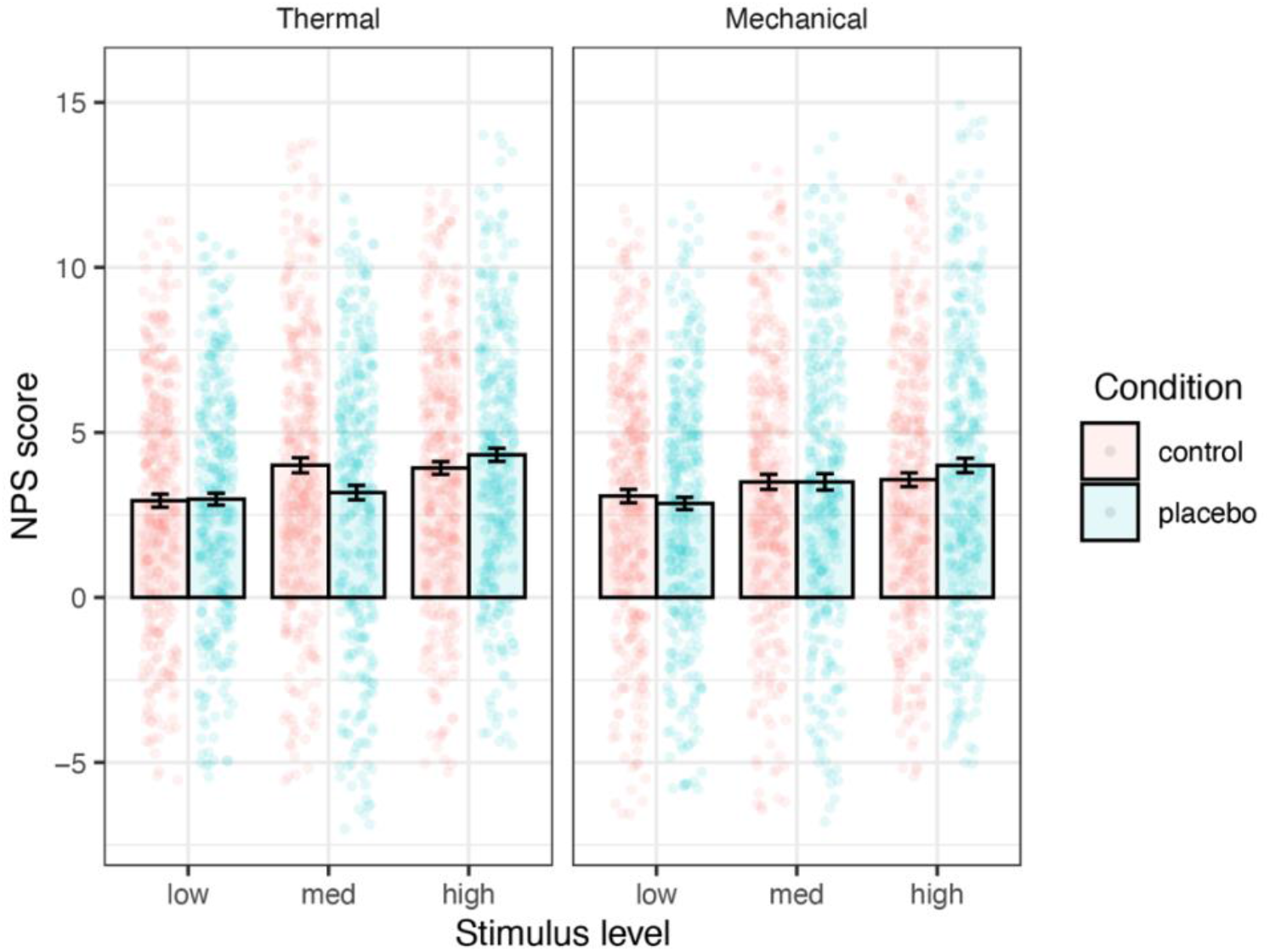
NPS results. The mean NPS scores (based on dot product) across participants are presented, in the Control (red) and Placebo (blue) condition, for each combination of modality and stimulus level. Error bars represent within-participant standard error of the mean, based on Morey, 2008 ^97^. Points represent single participants. Note that for better visualization, the 2.5% lowest and 2.5% highest observations from each category were excluded (the bars and error bars are based on the entire distribution). Asterisks represent significance of the placebo effect (Placebo vs. Control, uncorrected): * *p* < .05, ** *p* < .01, *** *p* < .001.

**Supplementary Figure 3.**
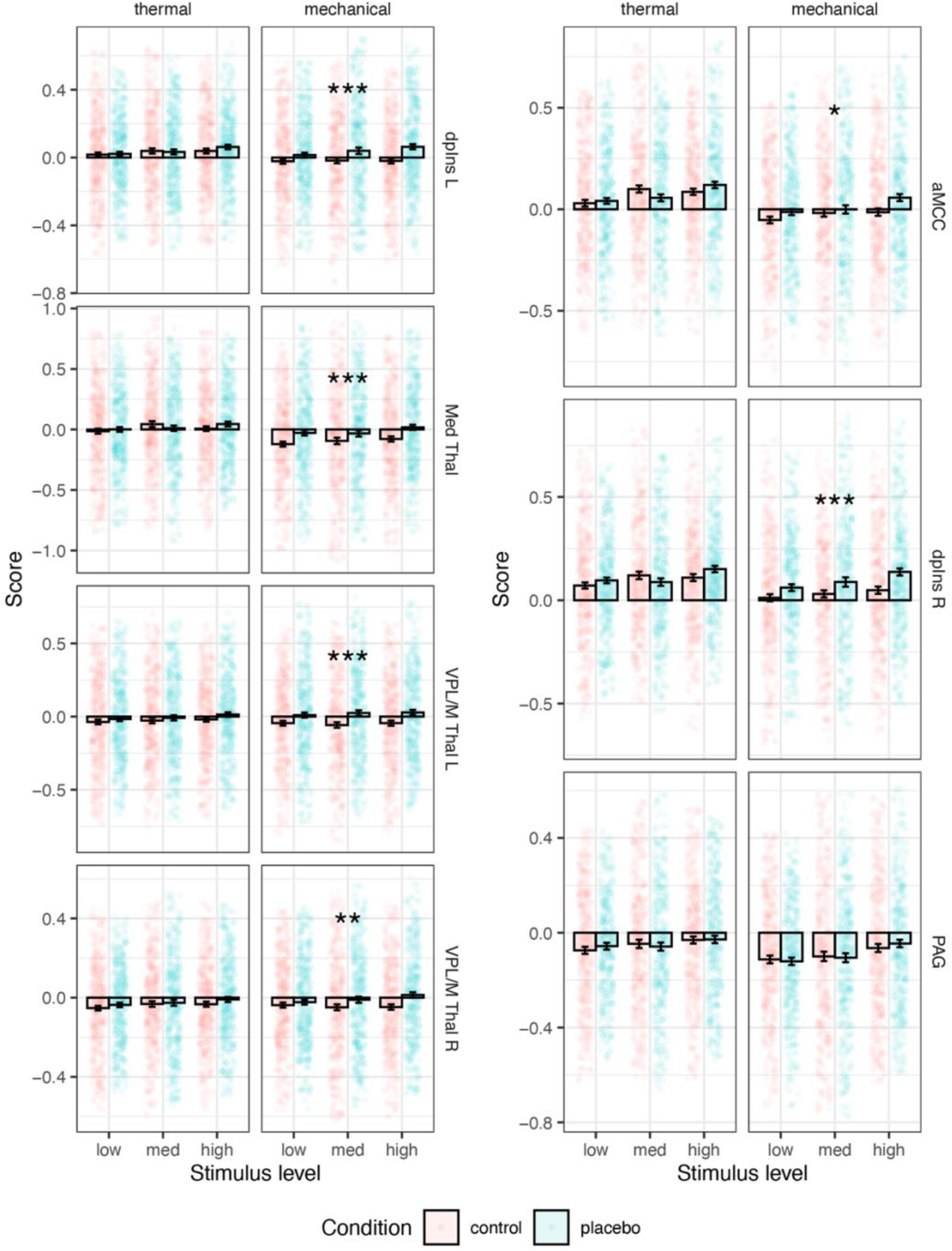
Activity in a priori nociceptive regions of interest. The mean activity across voxels in each a priori nociceptive region across participants is presented, in the Control (red) and Placebo (blue) condition, for each combination of modality and stimulus level. Error bars represent within-participant standard error of the mean, based on Morey, 2008 ^97^. Note that for better visualization, the 5% lowest and 5% highest observations from each category were excluded (the bars and error bars are based on the entire distribution). Abbreviations: L (left), R (right), aMCC (anterior midcingulate cortex), dpIns (dorsal posterior insula); Med Thal (medial thalamus); VPL/M Thal (ventral posterior thalamus). Asterisks represent significance of the placebo effect (Placebo vs. Control, uncorrected): * *p* < .05, ** *p* < .01, *** *p* < .001.

**Supplementary Figure 4.**
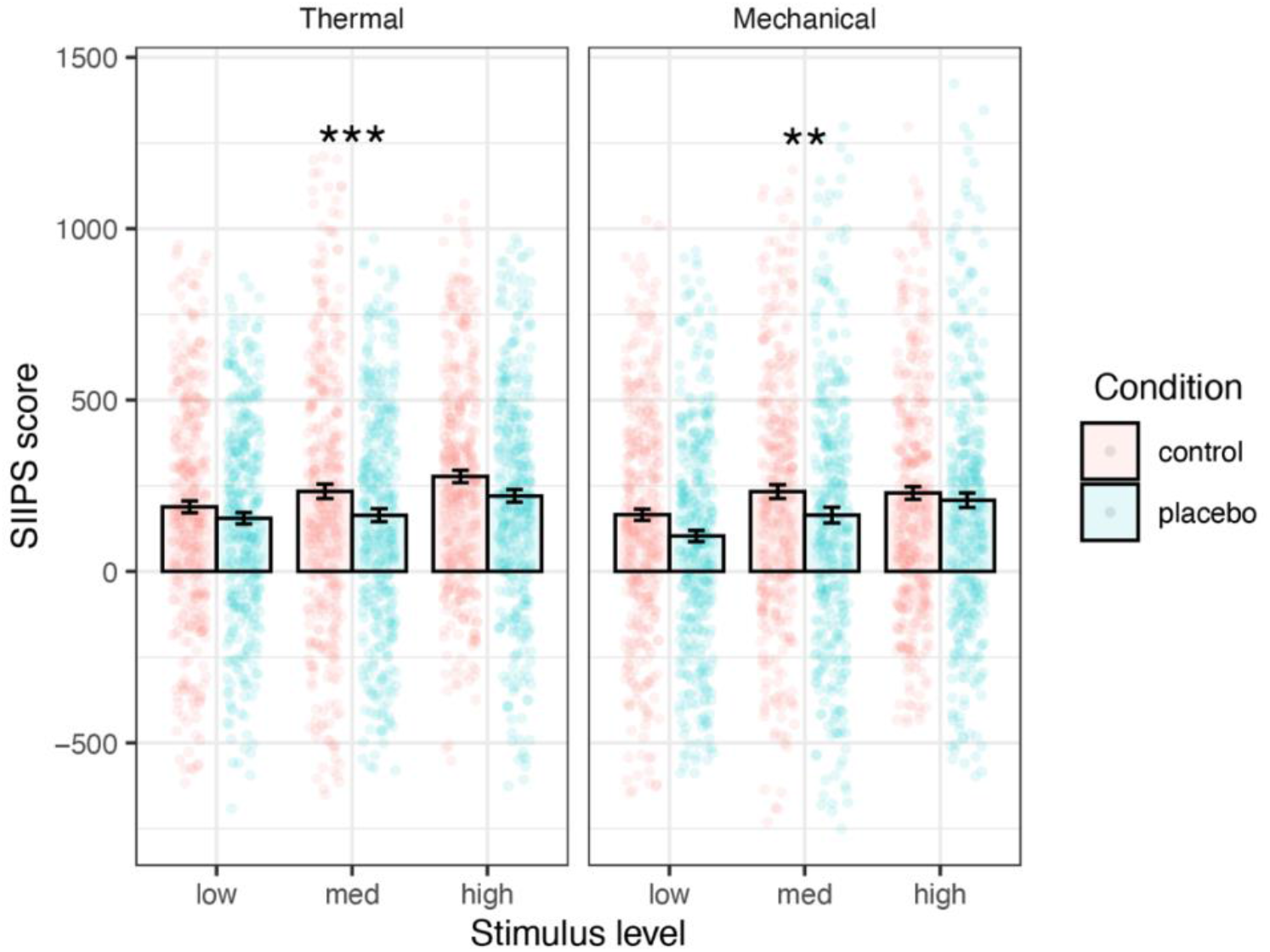
SIIPS results. The mean SIIPS scores (based on dot product) across participants are presented, in the Control (red) and Placebo (blue) condition, for each combination of modality and stimulus level. Error bars represent within-participant standard error of the mean, based on Morey, 2008 ^97^. Points represent single participants. Note that for better visualization, the 2.5% lowest and 2.5% highest observations from each category were excluded (the bars and error bars are based on the entire distribution). Asterisks represent significance of the placebo effect (Placebo vs. Control, uncorrected): * *p* < .05, ** *p* < .01, *** *p* < .001.

**Supplementary Figure 5.**
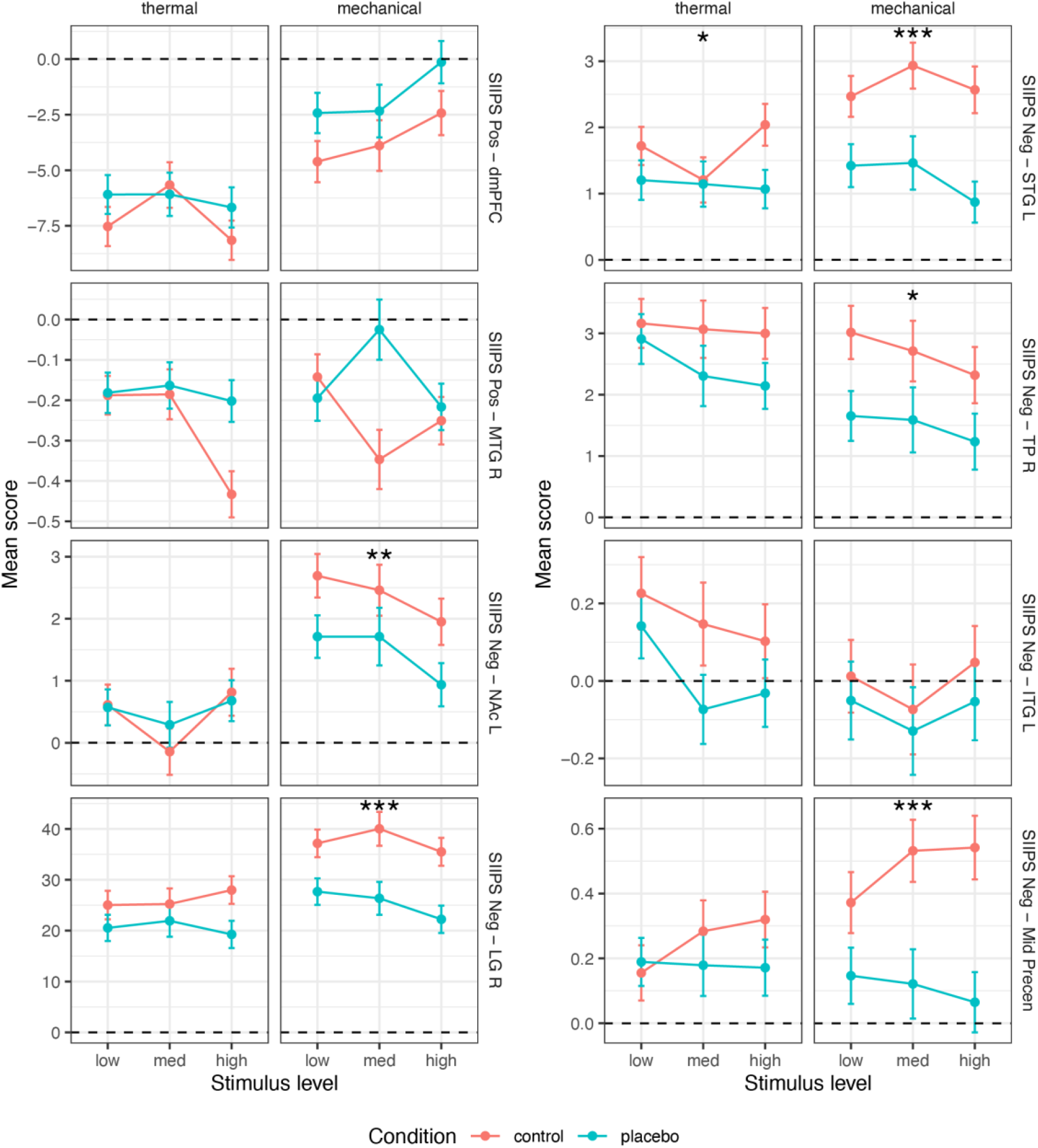
Activity in a priori SIIPS’ subregions. The local pattern responses of each a priori subregion of SIIPS across participants are presented, in the Control (red) and Placebo (blue) condition, for each combination of modality and stimulus level. Error bars represent within-participant standard error of the mean, based on Morey, 2008 ^97^. Asterisks represent significance of the placebo effect (Placebo vs. Control, ucorrected): * *p* < .05, ** *p* < .01, *** *p* < .001. See also Supplementary Figure 6. Abbreviation: SIIPS (Stimulus Intensity Independent Pain Signature); Pos (positive); Neg (negative); L (left); R (right); dmPFC (dorsomedial prefrontal cortex); MTG (middle temporal gyrus); NAc (nucleus accumbens); LG (lingual gyrus); STG (superior temporal gyrus); TP (temporal pole); ITG (inferior temporal gyrus); mid precen (middle precentral gyrus).

**Supplementary Figure 6.**
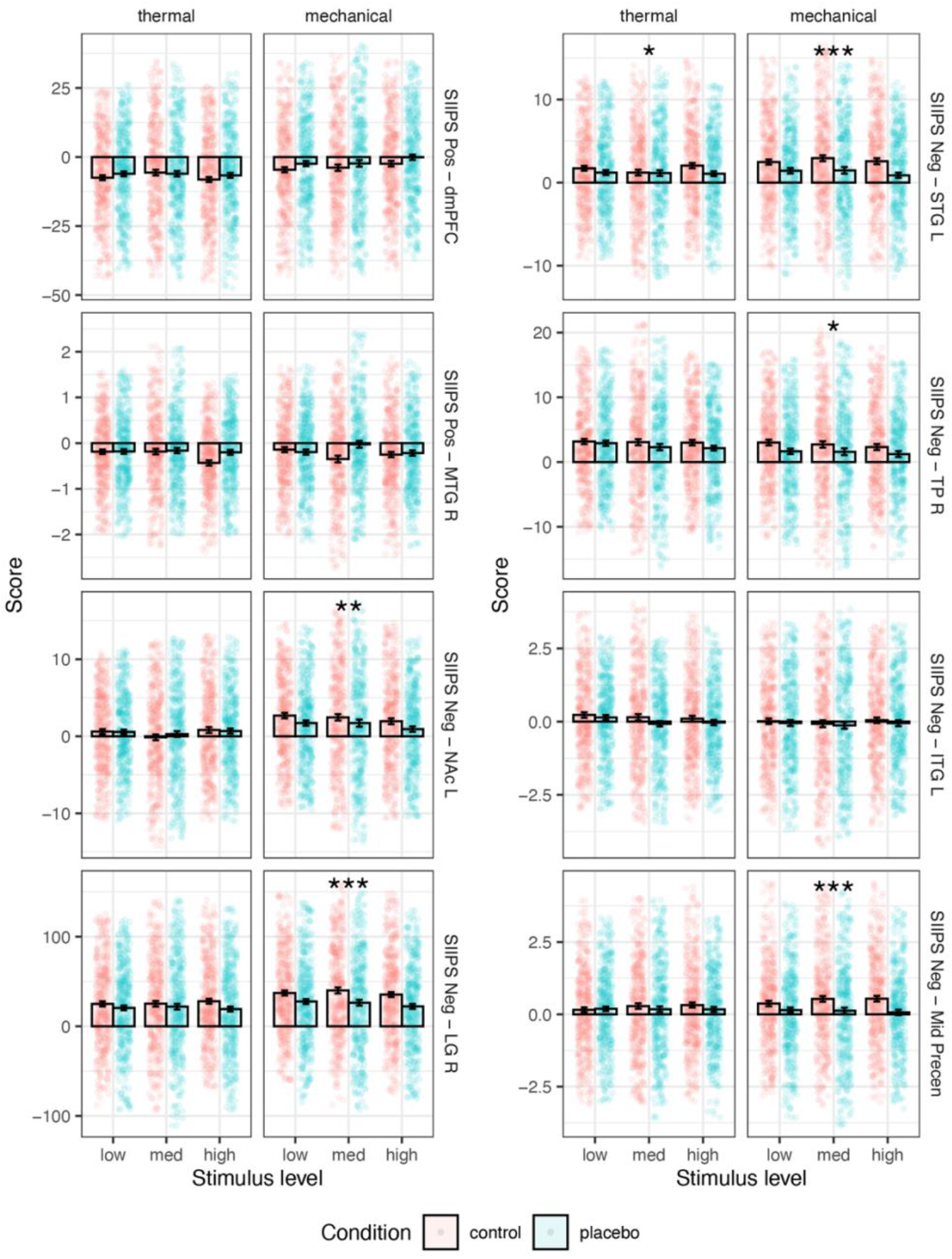
Activity in a priori SIIPS’ subregions. The local pattern responses of each a priori subregion across participants is presented, in the Control (red) and Placebo (blue) condition, for each combination of modality and stimulus level. Error bars represent within-participant standard error of the mean, based on Morey, 2008 ^97^. Points represent single participants. Note that for better visualization, the 5% lowest and 5% highest observations from each category were excluded (the bars and error bars are based on the entire distribution). Asterisks represent significance of the placebo effect (Placebo vs. Control, uncorrected): * *p* < .05, ** *p* < .01, *** *p* < .001. Abbreviations: SIIPS (Stimulus Intensity Independent Pain Signature); Pos (positive); Neg (negative); L (left); R (right); dmPFC (dorsomedial prefrontal cortex); MTG (middle temporal gyrus); NAc (nucleus accumbens); LG (lingual gyrus); STG (superior temporal gyrus); TP (temporal pole); ITG (inferior temporal gyrus); mid precen (middle precentral gyrus).

**Supplementary Figure 7.**
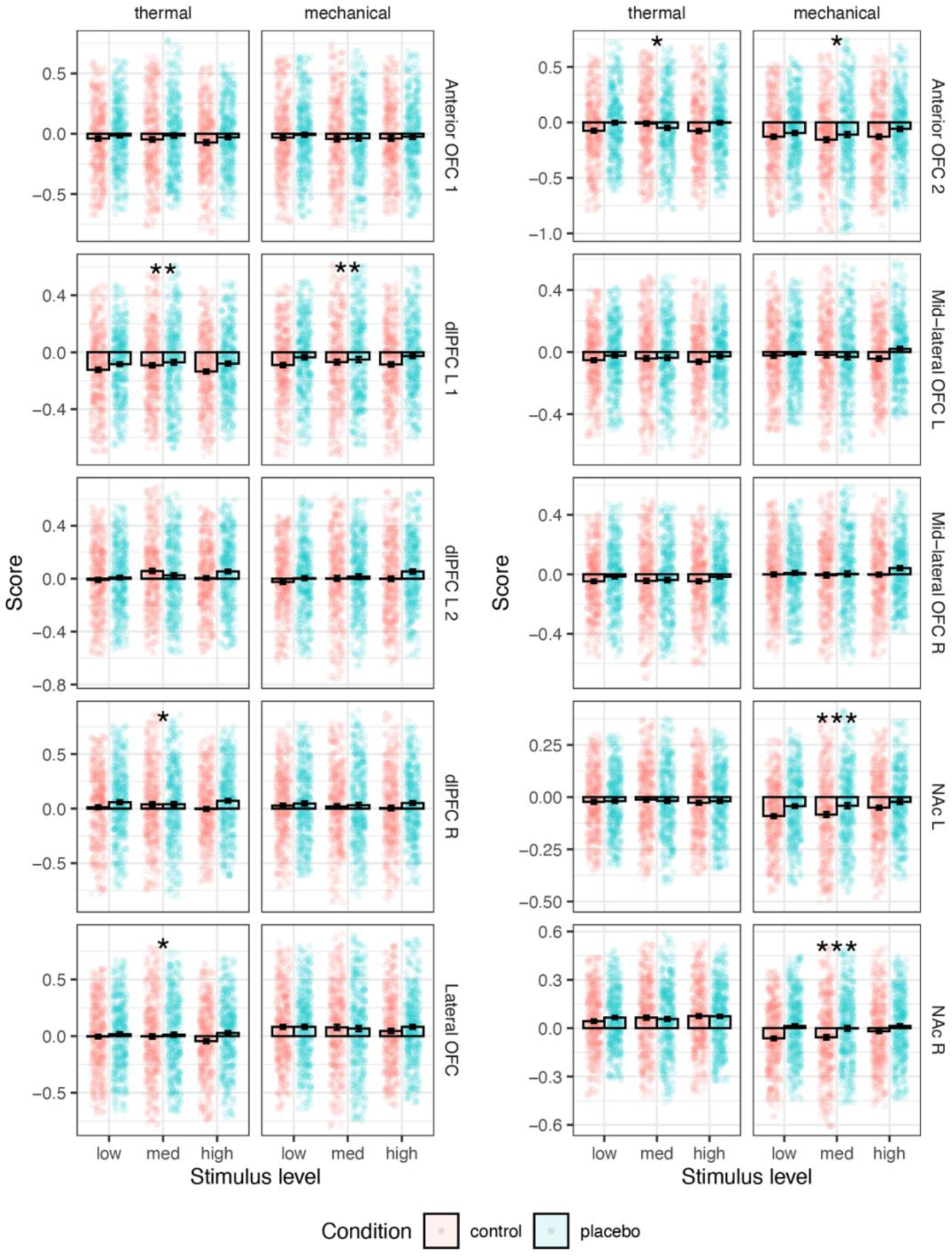
Activity in a priori higher-level pain processing regions. The mean activity across voxels in each a priori ROI across participants is presented, in the Control (red) and Placebo (blue) condition, for each combination of modality and stimulus level. Error bars represent within-participant standard error of the mean, based on Morey, 2008 ^97^. Points represent single participants. Note that for better visualization, the 5% lowest and 5% highest observations from each category were excluded (the bars and error bars are based on the entire distribution). Asterisks represent significance of the placebo effect (Placebo vs. Control, uncorrected): * *p* < .05, ** *p* < .01, *** *p* < .001. Abbreviations: L (left); R (right); NAc (nucleus accumbens); dlPFC (dorsolateral prefrontal cortex); OFC (orbitofrontal cortex).

## Supplementary Tables

**Supplementary Table 1.**
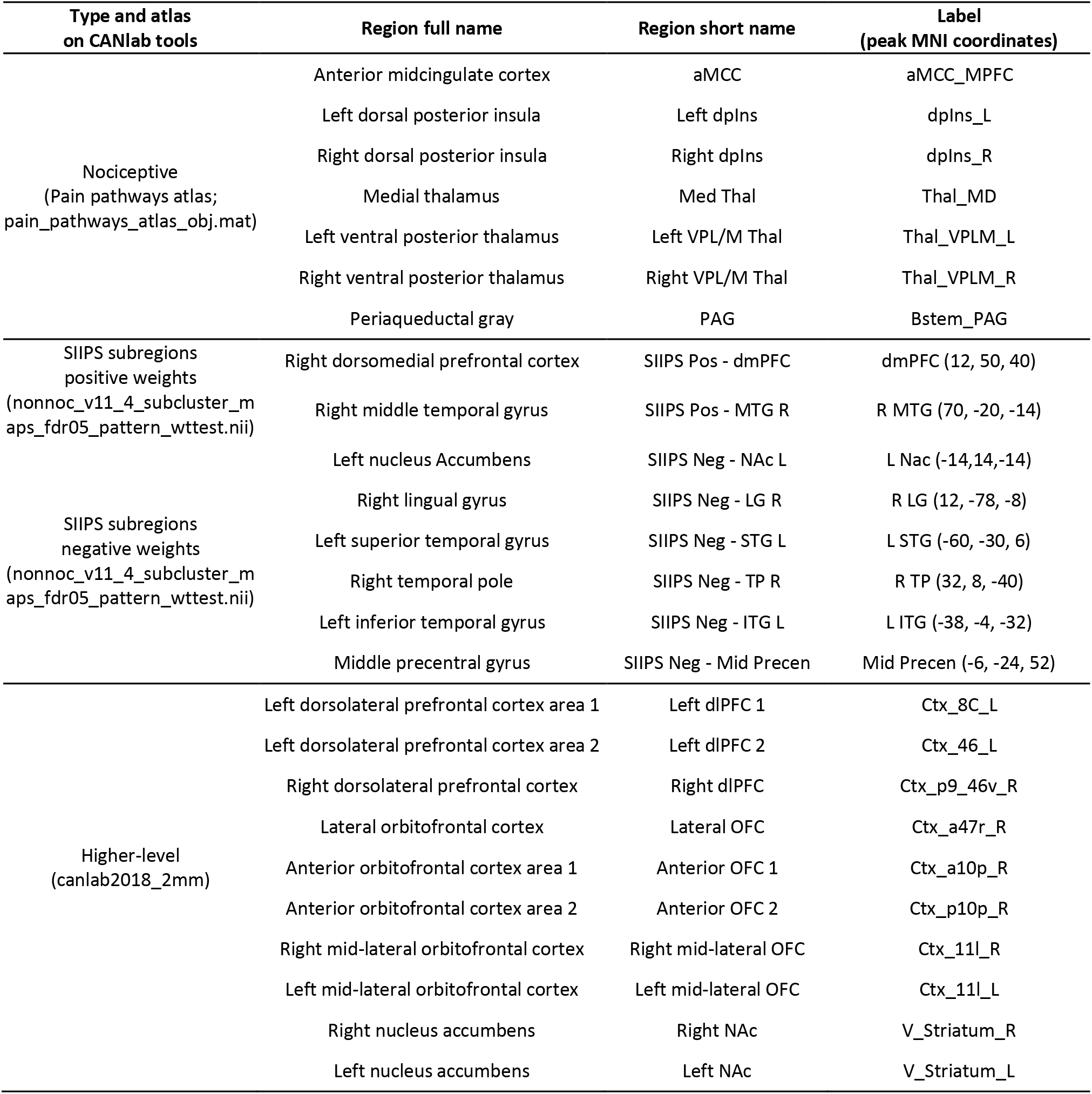
A priori pre-registered regions of interest.

**Supplementary Table 2.** Bayes Factors for a priori nociceptive neuromarker and ROIs.

|  |  | NPS | Left<br>dplns | Right<br>dplns | aMCC | Left VPL/M<br>Thal | Right VPL/M<br>Thal | Medial<br>Thal | PAG |
| --- | --- | --- | --- | --- | --- | --- | --- | --- | --- |
| Thermal | wide | 0.044<br>(0.098) | 0.037<br>(0.072) | 0.048<br>(0.060) | 0.034<br>(0.132) | 0.143<br>(0.190) | 0.086<br>(0.058) | 0.033<br>(0.134) | 0.033<br>(0.033) |
|  | wide with<br>interaction | 0.056<br>(0.097) | 0<br>(0.363) | 0.004<br>(0.054) | 0.004<br>(0.117) | 0.001<br>(0.334) | 0.001<br>(0.123) | 0.001<br>(0.136) | 0<br>(0.040) |
|  | regular<br>width | 0.069<br>(0.234) | 0.160<br>(0.641) | 0.082<br>(0.126) | 0.053<br>(0.215) | 0.126<br>(0.411) | 0.121<br>(0.076) | 0.048<br>(0.113) | 0.048<br>(0.032) |
|  | ultrawide | 0.036<br>(0.172) | 0.027<br>(0.082) | 0.034<br>(0.113) | 0.020<br>(0.099) | 0.102<br>(0.133) | 0.057<br>(0.076) | 0.025<br>(0.066) | 0.025<br>(0.069) |
| Mechanical | wide | 0.036<br>(0.117) | 820.7<br>(0.226) | 374.3<br>(0.234) | 2<br>(0.081) | 1125.3<br>(0.181) | 5.9<br>(0.091) | 394.4<br>(0.132) | 0.032<br>(0.069) |
|  | wide with<br>interaction | 0.001<br>(0.096) | 8.5<br>(0.112) | 3.1<br>(0.203) | 0.036<br>(0.077) | 7.5<br>(0.196) | 0.110<br>(0.072) | 2.547<br>(0.091) | 0<br>(0.069) |
|  | regular<br>width | 0.049<br>(0.071) | 785.2<br>(0.071) | 614.5<br>(0.378) | 2.9<br>(0.091) | 1674.3<br>(0.066) | 8.3<br>(0.089) | 412.2<br>(0.100) | 0.048<br>(0.058) |
|  | ultrawide | 0.024<br>(0.050) | 447.3<br>(0.134) | 213<br>(0.108) | 1.3<br>(0.087) | 631<br>(0.464) | 3.8<br>(0.083) | 289.8<br>(0.196) | 0.023<br>(0.057) |

Bayes Factor (BF) values for tests of placebo effect in a priori nociceptive neuromarker and ROIs. BF values (evidence in favor of the alternative hypothesis divided by the evidence in favor of the null hypothesis) are presented, along with the corresponding error of the BF value in parenthesis, for each neuromarker / ROI, modality, width of the prior distribution ( $\sqrt{2}/2$ , 1, and $\sqrt{2}$ ) and inclusion of the interaction term. Abbreviations: aMCC (anterior midcingulate cortex), dplns (dorsal posterior insula); thal (thalamus); VPL/M (ventral posterior thalamus); PAG (periaqueductal gray).

**Supplementary Table 3.**
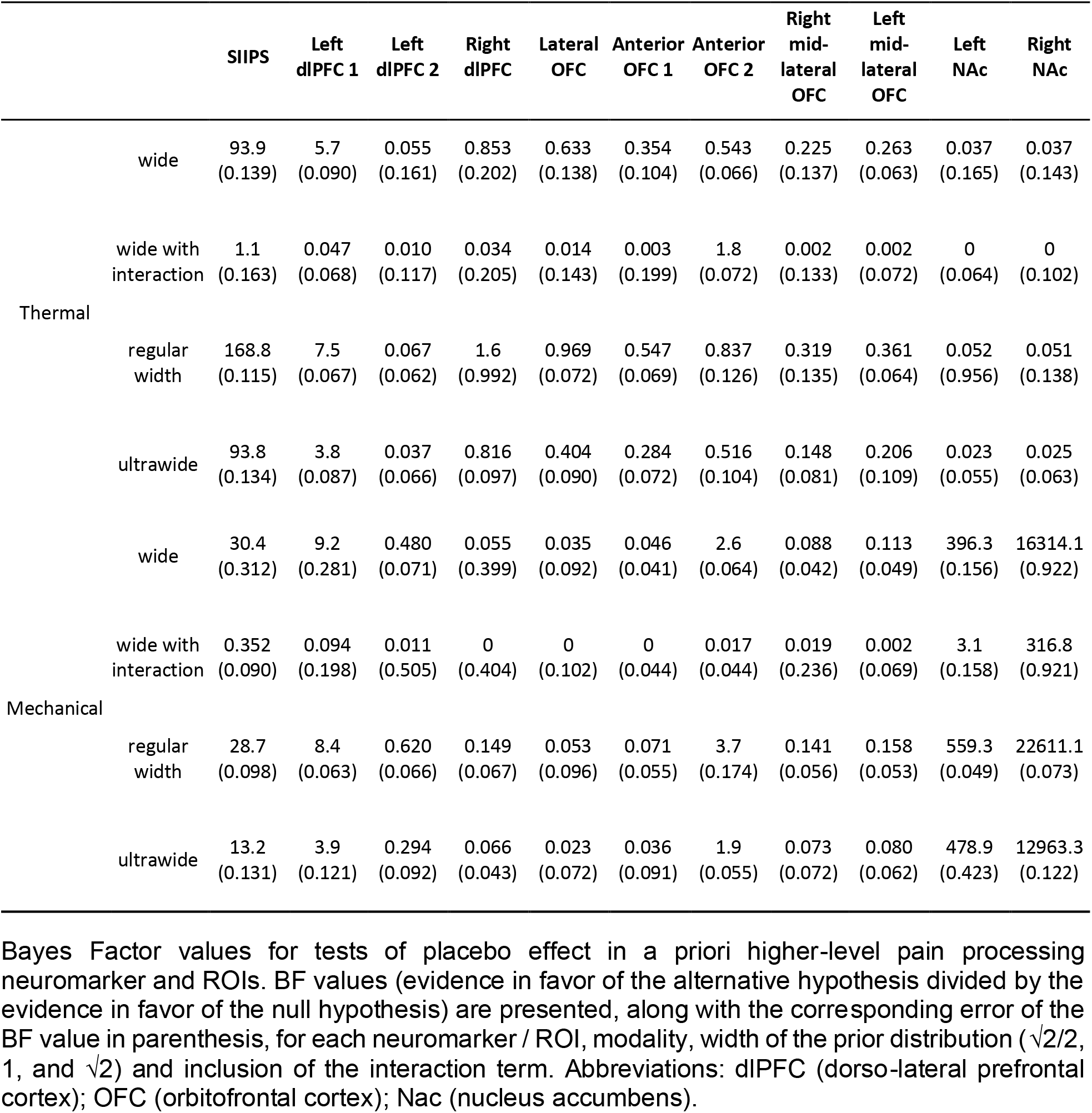
Bayes Factors for a priori higher-level pain processing neuromarker and ROIs.

**Supplementary Table 4.**
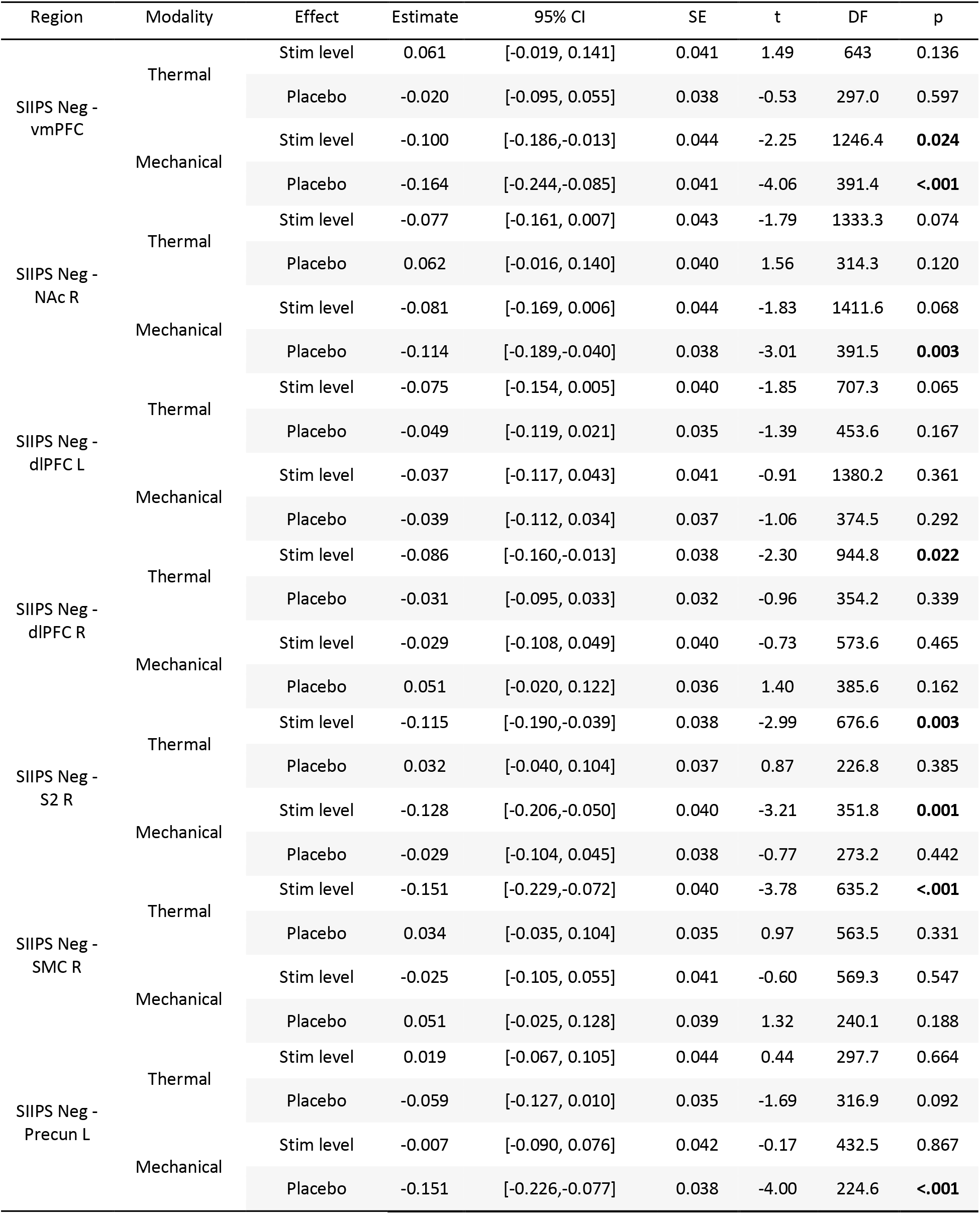
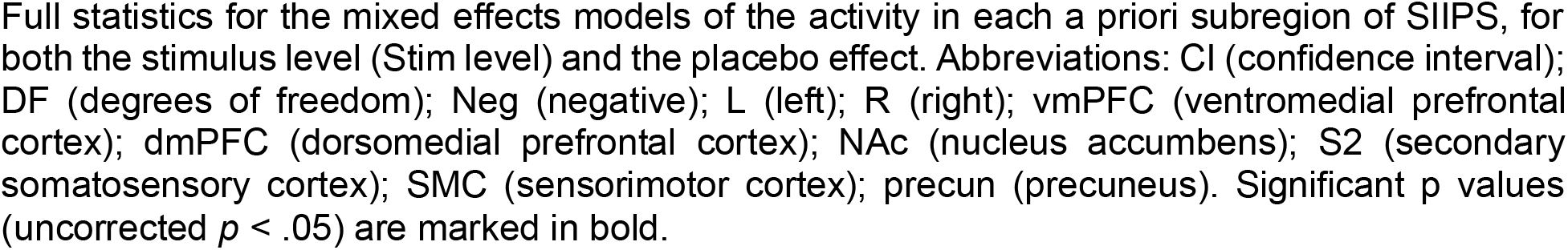
Statistics for additional, non-pre-registered subregions of SIIPS: stimulus level and placebo effects.

| Region | Modality | Effect | Estimate | 95% CI | SE | t | DF | p |
| --- | --- | --- | --- | --- | --- | --- | --- | --- |
| SIIPS Neg - vmPFC | Thermal | Stim level | 0.061 | [-0.019, 0.141] | 0.041 | 1.49 | 643 | 0.136 |
|  |  | Placebo | -0.020 | [-0.095, 0.055] | 0.038 | -0.53 | 297.0 | 0.597 |
|  | Mechanical | Stim level | -0.100 | [-0.186,-0.013] | 0.044 | -2.25 | 1246.4 | **0.024** |
|  |  | Placebo | -0.164 | [-0.244,-0.085] | 0.041 | -4.06 | 391.4 | **<.001** |
| SIIPS Neg - NAc R | Thermal | Stim level | -0.077 | [-0.161, 0.007] | 0.043 | -1.79 | 1333.3 | 0.074 |
|  |  | Placebo | 0.062 | [-0.016, 0.140] | 0.040 | 1.56 | 314.3 | 0.120 |
|  | Mechanical | Stim level | -0.081 | [-0.169, 0.006] | 0.044 | -1.83 | 1411.6 | 0.068 |
|  |  | Placebo | -0.114 | [-0.189,-0.040] | 0.038 | -3.01 | 391.5 | **0.003** |
| SIIPS Neg - dIPFC L | Thermal | Stim level | -0.075 | [-0.154, 0.005] | 0.040 | -1.85 | 707.3 | 0.065 |
|  |  | Placebo | -0.049 | [-0.119, 0.021] | 0.035 | -1.39 | 453.6 | 0.167 |
|  | Mechanical | Stim level | -0.037 | [-0.117, 0.043] | 0.041 | -0.91 | 1380.2 | 0.361 |
|  |  | Placebo | -0.039 | [-0.112, 0.034] | 0.037 | -1.06 | 374.5 | 0.292 |
| SIIPS Neg - dIPFC R | Thermal | Stim level | -0.086 | [-0.160,-0.013] | 0.038 | -2.30 | 944.8 | **0.022** |
|  |  | Placebo | -0.031 | [-0.095, 0.033] | 0.032 | -0.96 | 354.2 | 0.339 |
|  | Mechanical | Stim level | -0.029 | [-0.108, 0.049] | 0.040 | -0.73 | 573.6 | 0.465 |
|  |  | Placebo | 0.051 | [-0.020, 0.122] | 0.036 | 1.40 | 385.6 | 0.162 |
| SIIPS Neg - S2 R | Thermal | Stim level | -0.115 | [-0.190,-0.039] | 0.038 | -2.99 | 676.6 | **0.003** |
|  |  | Placebo | 0.032 | [-0.040, 0.104] | 0.037 | 0.87 | 226.8 | 0.385 |
|  | Mechanical | Stim level | -0.128 | [-0.206,-0.050] | 0.040 | -3.21 | 351.8 | **0.001** |
|  |  | Placebo | -0.029 | [-0.104, 0.045] | 0.038 | -0.77 | 273.2 | 0.442 |
| SIIPS Neg - SMC R | Thermal | Stim level | -0.151 | [-0.229,-0.072] | 0.040 | -3.78 | 635.2 | **<.001** |
|  |  | Placebo | 0.034 | [-0.035, 0.104] | 0.035 | 0.97 | 563.5 | 0.331 |
|  | Mechanical | Stim level | -0.025 | [-0.105, 0.055] | 0.041 | -0.60 | 569.3 | 0.547 |
|  |  | Placebo | 0.051 | [-0.025, 0.128] | 0.039 | 1.32 | 240.1 | 0.188 |
| SIIPS Neg - Precun L | Thermal | Stim level | 0.019 | [-0.067, 0.105] | 0.044 | 0.44 | 297.7 | 0.664 |
|  |  | Placebo | -0.059 | [-0.127, 0.010] | 0.035 | -1.69 | 316.9 | 0.092 |
|  | Mechanical | Stim level | -0.007 | [-0.090, 0.076] | 0.042 | -0.17 | 432.5 | 0.867 |
|  |  | Placebo | -0.151 | [-0.226,-0.077] | 0.038 | -4.00 | 224.6 | **<.001** |

Full statistics for the mixed effects models of the activity in each a priori subregion of SIIPS, for both the stimulus level (Stim level) and the placebo effect. Abbreviations: CI (confidence interval); DF (degrees of freedom); Neg (negative); L (left); R (right); vmPFC (ventromedial prefrontal cortex); dmPFC (dorsomedial prefrontal cortex); NAc (nucleus accumbens); S2 (secondary somatosensory cortex); SMC (sensorimotor cortex); precun (precuneus). Significant $p$ values (uncorrected $p < .05$ ) are marked in bold.

**Supplementary Table 5.** Statistics for a priori nociceptive ROIs: correlation between behavioral and neural placebo-induced reductions.

| Region | Modality | Estimate | 95% CI | SE | t | DF | p |
| --- | --- | --- | --- | --- | --- | --- | --- |
| aMCC | Thermal | 0.152 | [0.084, 0.220] | 0.034 | 4.47 | 88.0 | **< .001** |
|  | Mechanical | 0.171 | [0.093, 0.248] | 0.039 | 4.38 | 94.3 | **< .001** |
| dplns L | Thermal | 0.056 | [-0.012, 0.124] | 0.034 | 1.63 | 129.9 | 0.106 |
|  | Mechanical | 0.142 | [0.068, 0.215] | 0.037 | 3.84 | 87.3 | **< .001** |
| dplns R | Thermal | 0.093 | [0.029, 0.157] | 0.033 | 2.86 | 230.5 | **0.005** |
|  | Mechanical | 0.136 | [0.064, 0.208] | 0.036 | 3.76 | 79.6 | **< .001** |
| PAG | Thermal | 0.100 | [0.033, 0.167] | 0.034 | 2.95 | 134.5 | **0.004** |
|  | Mechanical | 0.167 | [0.096, 0.239] | 0.036 | 4.70 | 49.6 | **< .001** |
| Med Thal | Thermal | 0.083 | [0.015, 0.151] | 0.035 | 2.41 | 180.2 | **0.017** |
|  | Mechanical | 0.065 | [-0.006, 0.136] | 0.036 | 1.81 | 219.3 | 0.071 |
| VPL/M Thal L | Thermal | 0.035 | [-0.029, 0.099] | 0.033 | 1.07 | 232.6 | 0.286 |
|  | Mechanical | 0.051 | [-0.023, 0.124] | 0.037 | 1.36 | 127.9 | 0.178 |
| VPL/M Thal R | Thermal | 0.035 | [-0.033, 0.103] | 0.034 | 1.02 | 117.2 | 0.311 |
|  | Mechanical | 0.054 | [-0.013, 0.121] | 0.034 | 1.58 | 144.1 | 0.116 |

Full statistics for the mixed effects models of the correlation of the Control - Placebo neural activity in each a priori ROI and the behavioral analgesia (Control - Placebo pain ratings). Abbreviations: CI (confidence interval); DF (degrees of freedom); L (left); R (right); aMCC (anterior midcingulate cortex); dplns (dorsal posterior insula); Med Thal (medial thalamus); VPL/M Thal (ventral posterior thalamus); PAG (periaqueductal gray). Significant p values (uncorrected $p < .05$ ) are marked in bold.

**Supplementary Table 6.** Statistics for a priori subregions of SIIPS: correlation between behavioral and neural placebo-induced reductions.

| Region | Modality | Estimate | 95% CI | SE | t | DF | p |
| --- | --- | --- | --- | --- | --- | --- | --- |
| SIIPS Pos - dmPFC | Thermal | 0.023 | [-0.042, 0.087] | 0.033 | 0.70 | 272.7 | 0.485 |
|  | Mechanical | 0.072 | [-0.004, 0.148] | 0.038 | 1.88 | 89 | 0.064 |
| SIIPS Pos - MTG R | Thermal | -0.020 | [-0.087, 0.047] | 0.034 | -0.59 | 226.3 | 0.555 |
|  | Mechanical | -0.005 | [-0.072, 0.061] | 0.034 | -0.16 | 171.3 | 0.873 |
| SIIPS Neg - NAc L | Thermal | -0.041 | [-0.105, 0.022] | 0.032 | -1.28 | 189.5 | 0.203 |
|  | Mechanical | -0.030 | [-0.104, 0.045] | 0.037 | -0.79 | 71.2 | 0.430 |
| SIIPS Neg - LG R | Thermal | 0.093 | [0.029, 0.157] | 0.032 | 2.88 | 199.5 | **0.004** |
|  | Mechanical | -0.011 | [-0.081, 0.058] | 0.035 | -0.32 | 106.5 | 0.747 |
| SIIPS Neg - STG L | Thermal | 0.011 | [-0.054, 0.076] | 0.033 | 0.33 | 128.8 | 0.743 |
|  | Mechanical | -0.091 | [-0.158, -0.024] | 0.034 | -2.70 | 120 | **0.008** |
| SIIPS Neg - TP R | Thermal | 0.023 | [-0.047, 0.093] | 0.035 | 0.65 | 146 | 0.519 |
|  | Mechanical | -0.079 | [-0.144, -0.013] | 0.033 | -2.35 | 218.3 | **0.020** |
| SIIPS Neg - ITG L | Thermal | 0.035 | [-0.029, 0.098] | 0.032 | 1.08 | 391.6 | 0.282 |
|  | Mechanical | -0.022 | [-0.088, 0.044] | 0.033 | -0.66 | 199.9 | 0.508 |
| SIIPS Neg - Mid Precen | Thermal | 0.029 | [-0.034, 0.091] | 0.032 | 0.90 | 354.4 | 0.368 |
|  | Mechanical | -0.028 | [-0.098, 0.043] | 0.035 | -0.78 | 90.2 | 0.438 |

Full statistics for the mixed effects models of the correlation of the Control - Placebo neural activity in each a priori subregion and the behavioral analgesia (Control - Placebo pain ratings). Abbreviations: CI (confidence interval); DF (degrees of freedom); Pos (positive); Neg (negative); L (left); R (right); dmPFC (dorsomedial prefrontal cortex); MTG (middle temporal gyrus); NAc (nucleus accumbens); LG (lingual gyrus); STG (superior temporal gyrus); TP (temporal pole); ITG (inferior temporal gyrus); mid precen (middle precentral gyrus). Significant p values (uncorrected $p < .05$ ) are marked in bold.

**Supplementary Table 7.**
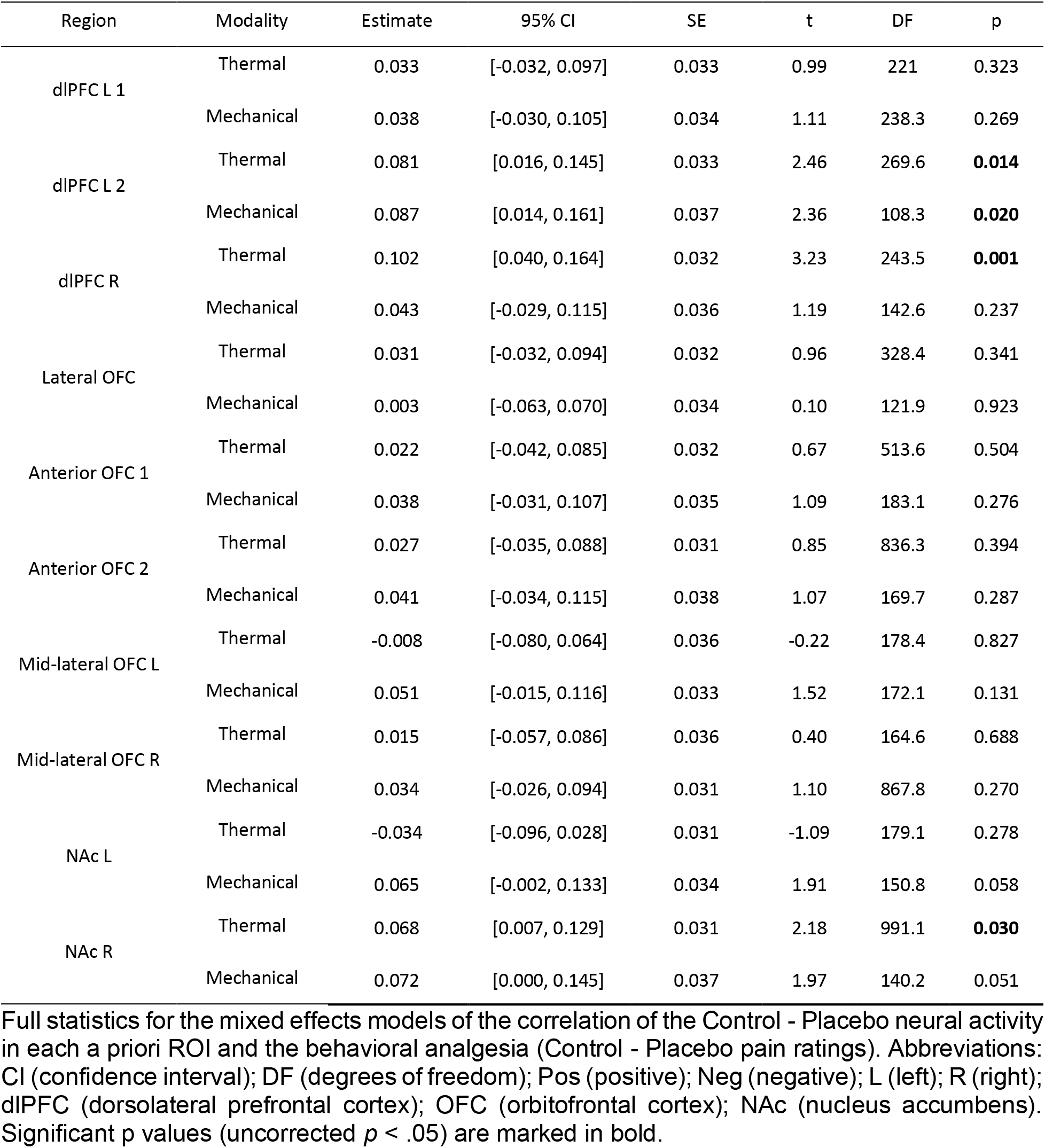
Statistics for a priori ROIs for higher-level pain processing: correlation between behavioral and neural placebo-induced reductions.

| Region | Modality | Estimate | 95% CI | SE | t | DF | p |
| --- | --- | --- | --- | --- | --- | --- | --- |
| dIPFC L 1 | Thermal | 0.033 | [-0.032, 0.097] | 0.033 | 0.99 | 221 | 0.323 |
|  | Mechanical | 0.038 | [-0.030, 0.105] | 0.034 | 1.11 | 238.3 | 0.269 |
| dIPFC L 2 | Thermal | 0.081 | [0.016, 0.145] | 0.033 | 2.46 | 269.6 | **0.014** |
|  | Mechanical | 0.087 | [0.014, 0.161] | 0.037 | 2.36 | 108.3 | **0.020** |
| dIPFC R | Thermal | 0.102 | [0.040, 0.164] | 0.032 | 3.23 | 243.5 | **0.001** |
|  | Mechanical | 0.043 | [-0.029, 0.115] | 0.036 | 1.19 | 142.6 | 0.237 |
| Lateral OFC | Thermal | 0.031 | [-0.032, 0.094] | 0.032 | 0.96 | 328.4 | 0.341 |
|  | Mechanical | 0.003 | [-0.063, 0.070] | 0.034 | 0.10 | 121.9 | 0.923 |
| Anterior OFC 1 | Thermal | 0.022 | [-0.042, 0.085] | 0.032 | 0.67 | 513.6 | 0.504 |
|  | Mechanical | 0.038 | [-0.031, 0.107] | 0.035 | 1.09 | 183.1 | 0.276 |
| Anterior OFC 2 | Thermal | 0.027 | [-0.035, 0.088] | 0.031 | 0.85 | 836.3 | 0.394 |
|  | Mechanical | 0.041 | [-0.034, 0.115] | 0.038 | 1.07 | 169.7 | 0.287 |
| Mid-lateral OFC L | Thermal | -0.008 | [-0.080, 0.064] | 0.036 | -0.22 | 178.4 | 0.827 |
|  | Mechanical | 0.051 | [-0.015, 0.116] | 0.033 | 1.52 | 172.1 | 0.131 |
| Mid-lateral OFC R | Thermal | 0.015 | [-0.057, 0.086] | 0.036 | 0.40 | 164.6 | 0.688 |
|  | Mechanical | 0.034 | [-0.026, 0.094] | 0.031 | 1.10 | 867.8 | 0.270 |
| NAc L | Thermal | -0.034 | [-0.096, 0.028] | 0.031 | -1.09 | 179.1 | 0.278 |
|  | Mechanical | 0.065 | [-0.002, 0.133] | 0.034 | 1.91 | 150.8 | 0.058 |
| NAc R | Thermal | 0.068 | [0.007, 0.129] | 0.031 | 2.18 | 991.1 | **0.030** |
|  | Mechanical | 0.072 | [0.000, 0.145] | 0.037 | 1.97 | 140.2 | 0.051 |

Full statistics for the mixed effects models of the correlation of the Control - Placebo neural activity in each a priori ROI and the behavioral analgesia (Control - Placebo pain ratings). Abbreviations: CI (confidence interval); DF (degrees of freedom); Pos (positive); Neg (negative); L (left); R (right); dIPFC (dorsolateral prefrontal cortex); OFC (orbitofrontal cortex); NAc (nucleus accumbens). Significant p values (uncorrected $p < .05$ ) are marked in bold.

